# Decoding Cortical Precursors of Evoked and Spontaneous Pupil Dilation

**DOI:** 10.64898/2026.05.27.726399

**Authors:** Takahito Komori, Shota Mizusaki, Shusuke Tomita, Nobuhiro Nakai, Toru Takumi, Naru Yoneda, Osamu Matoba, Mitsuhiro Morita

## Abstract

Pupil dilation reflects arousal and occurs both in response to sensory events and spontaneously. However, whether these two forms of dilation are preceded by distinct cortical dynamics remains unclear. Here, we combined pupillometry, wide-field Ca²⁺ imaging in mice, and explainable machine learning to estimate cortical activity patterns that predict pupil dilation. Evoked dilation was examined during somatosensory, visual pattern, and visual context stimulations, and spontaneous dilation was analyzed using a public resting-state dataset. Independent components of cortical activity were used as inputs to recurrent neural network (RNN) decoders. When sessions with pre-existing dilation were excluded, the decoders predicted evoked dilation above that of the shuffled-label controls. SHapley Additive exPlanations (SHAP) with spatial back-mapping revealed that activity around the retrosplenial cortex contributed to predictions across sensory modalities, whereas secondary motor cortex activity was prominent specifically during visual stimulation. In contrast, spontaneous dilation required lower-frequency cortical signals for decoding and showed a reverse order, with medial anterior activity preceding retrosplenial and secondary motor contributions. These findings indicate that evoked and spontaneous pupil dilation are preceded by distinct cortical dynamics, with evoked dilation involving a retrosplenial signal shared across sensory modalities and subsequent recruitment of the secondary motor cortex during visual stimulation.

## Introduction

Pupil diameter covaries with noradrenergic locus coeruleus (LC) activity and has long been used as an index of arousal state (Hall and Chilcott 2018; Privitera et al. 2020). The pupil dilates both spontaneously and in response to sensory stimulation, a phenomenon linked to attention, prediction error, novelty, cognitive load, exploratory behavior, and emotion (Bradley et al. 2008; Gilzenrat et al. 2010; Preuschoff et al. 2011; Nassar et al. 2012; Alnæs et al. 2014; Liao et al. 2016; Urai et al. 2017). Pupil dilation and the associated neural activity are altered in cognitive and psychiatric disorders (de Vries et al. 2021; Pelegrino et al. 2023; David et al. 2025). Characterizing the spatiotemporal distribution of cortical activity linked to pupil dilation is therefore important for understanding the neural basis of cognition and perception.

Resting-state pupil dilation is associated with tonic LC activation, which is linked to stress-induced amygdala activation and the default mode network (Aston-Jones and Cohen 2005; Rodriguez-Romaguera et al. 2020), and the arousal states reflected in pupil diameter are coupled with infra-slow cortical activity (Takahashi et al. 2023; Brink et al. 2016; Vinck et al. 2015; Carter et al. 2010; Raut et al. 2025). These findings largely emerged from pupillometry combined with fMRI and EEG. Wide-field Ca²⁺ imaging now measures mouse cortical activity at higher spatiotemporal resolution (Raut et al. 2025; Musall et al. 2023; Couto et al. 2021; Makino et al. 2017; Musall et al. 2019; Nakai et al. 2023). Combined with pupillometry and diverse sensory stimulation, it offers an opportunity to characterize the cortical activity accompanying pupil dilation in greater detail.

However, such data are high-dimensional and difficult to interpret. Conventional analyses, including inter-regional Pearson correlation based on the Allen Common Coordinate Framework (CCF) (Wang et al. 2020), principal component analysis (PCA), and localized semi-nonnegative matrix factorization (LocaNMF) (Saxena et al. 2020), identify activity correlated with pupil dilation, but not which patterns precede and thereby predict it. Machine learning has been applied successfully to decode behavioral states and pathological conditions from neural activity based on learned feature importance (Heinsfeld et al. 2018; Mitelut et al. 2022; Zhang et al. 2022; Ajioka et al. 2024; Raut et al. 2025).

Therefore, we combined wide-field Ca²⁺ imaging with machine learning-based analysis to investigate the cortical activity preceding pupil dilation, hypothesizing that sensory-evoked dilation is preceded by specific patterns of cortical activity involved in sensory perception. Models were trained to predict pupil state from cortical Ca²⁺ signals, and the spatiotemporal profile of the activity used for prediction was taken as an indicator of sensory processing leading to dilation. We compared three stimulation paradigms—electrical hindpaw, visual pattern, and visual context stimulation—and examined spontaneous dilation as a reference using a publicly available dataset (Kondo et al. 2025). Cortical activity shared across sensory modalities may constitute a core pathway linking sensory stimuli to perception and cognition, providing a starting point for circuit-level dissection.

## Materials and Methods

### Animal treatment

All animal experiments were approved by the Institutional Animal Care and Use Committee of Kobe University (approval number: 2021-03-01-R1). Three mice expressing GCaMP6f in excitatory cortical neurons (Emx1-Cre/Rosa-GCaMP6f) (Gorski et al. 2002; Madisen et al. 2015) (one male, two females; ≥6 weeks of age; 20–30 g) were used. Under ketamine–xylazine anesthesia, the skull was exposed and treated with 4-META/MMA-TBB dental resin for transcranial optical clearing, and a light-shielding frame and an aluminum headplate were secured to it (Supplementary Methods). Mice were housed individually for at least seven days and habituated to head fixation for at least three days without stimulus presentation.

### Wide-field Ca²⁺ imaging

All imaging was performed in awake mice, head-fixed under a custom-built fluorescence macroscope, with stimulus delivery and image acquisition synchronized via TTL signals. Each 40 s recording constituted one session; an experiment comprised 11 sessions (somatosensory and visual pattern stimulation) or eight sessions (visual context stimulation), initiated every 180 s. The first session of each experiment was excluded to avoid the initial transient response to LED onset.

GCaMP6f was excited alternately at 480 nm and 405 nm (20 ms each with a 5 ms interval; 40 frames/s in total), and fluorescence was detected by a scientific CMOS camera at binning 4x4 and sub-arrayed for 256 × 270 pixels. To image the full extent of the dorsal cortex, fluorescence was projected onto the sensor via a 2× objective lens and a 0.7× tube lens, yielding a spatial resolution of 0.0371 mm/pixel. Each pair of consecutive 405 nm and 480 nm frames was combined into one hemodynamic-corrected image (20 Hz). Images were registered to a cortical parcellation atlas (Supplementary Fig. 1) based on version 3 of the Allen Mouse Brain Common Coordinate Framework (CCF v3) (Wang et al. 2020) and converted to ΔF/F. Hemodynamic correction regressed the 405 nm signal out of the 480 nm signal within a 34-component singular value decomposition (SVD) subspace (Supplementary Methods).

### Pupillometry

The right eye was illuminated with a near-infrared lamp and recorded at 60 Hz with an infrared camera controlled by a Raspberry Pi, which was initiated by the same TTL signals that triggered image acquisition, synchronizing the onset of recording rather than individual frames. Pupil images were downsampled to 20 Hz, eye and pupil coordinates were extracted using DeepLabCut (Mathis et al. 2018), and horizontal pupil diameter was computed as the Euclidean distance between the left and right pupil boundary landmarks. Outliers identified by the sliding-window criterion and visual inspection were replaced with the preceding frame’s value (Supplementary Methods).

### Somatosensory stimulation

The body was gently secured to a metal plate, and two ECG electrodes were affixed to the plantar surface of the right hindpaw, approximately 1 mm apart. Transcutaneous electrical stimulation (pulse duration, 500 ms as in a previous study (Makino et al. 2017)) was delivered 20 s after the start of image acquisition at the threshold intensity (0.08–0.12 mA across animals), defined as the minimum intensity producing pupil dilation in at least three of five trials (Supplementary Methods).

### Visual stimulation

Head-fixed mice were placed on a custom treadmill to permit locomotion (Fig. 2A, left) with a display 60° to the left of the midline and a mirror symmetrically on the right, approximately 10 cm from the mouse. Sinusoidal gratings (0.1 cycles/degree) drifted at 3 cycles/s. No reward or aversive conditioning was applied. In the visual pattern experiment, gratings drifted continuously at one of four orientations (0°, 45°, 90°, or 135°); the stimulus event was a 90° clockwise shift in drift direction, maintained for 10 s and given 20 s after session onset (Fig. 2A, right). Gratings continued to drift between sessions, and direction was maintained throughout the experiment.

The visual context experiment employed an oddball paradigm (Bastos et al. 2012; Stefanics et al. 2014). Each presentation consisted of a 0.5 s halt (Halt) and 0.5 s drift (Drift), and eight consecutive presentations constituted an 8 s block (Fig. 3A). Drift direction was randomized in control blocks (Ctrl), whereas in test blocks (Test) it was constant during presentations 1–5 and 7–8 (redundant) and rotated 90° clockwise during presentation 6 (deviant). Each 40 s session comprised two Ctrl and three test blocks (Fig. 3A; Supplementary Methods).

### Resting state analysis using open-source dataset

Resting-state analyses used all data in an open-source dataset (DANDI: https://dandiarchive.org/dandiset/001425/) (Kondo et al. 2025), which includes simultaneous wide-field Ca²⁺ imaging and pupillometry from 25 vGluT1-Cre × Ai162 mice acquired at 30 Hz. Images were spatially downsampled and non-cortical pixels were masked to yield 256 × 270 pixel images matching the sensory-evoked analyses; the original sampling rate was retained.

### Pupil labels and target time window

Pupil diameter was low-pass filtered at 2 Hz (somatosensory) or 0.2 Hz (visual and resting-state experiments), Z-scored, min-max normalized, and binarized at 0.6 to create binary dilation labels. The pupil diameter peak was the maximum of the normalized trace in each trial, occurring 0.5 s or later after stimulus onset. To avoid under-or overestimation of positive labels (dilation events), the target time window relative to the peak was set from −0.9 to +0.5 s (somatosensory), −1.5 to +0.5 s (visual pattern), or −1.2 to +0.5 s (visual context). Resting-state recordings were segmented into 40 s epochs and labeled in the same way over a window from −3 to +0.5 s (Supplementary Methods).

### Cortical features

Spatially independent cortical components were extracted using the JADE ICA algorithm (Hyvärinen et al. 2001): for each condition, every fifth session was used to estimate the decomposition matrix from the top 200 SVD temporal components (cumulative explained variance > 99.5%). ICs with spatially coherent, anatomically interpretable patterns were selected by visual inspection. The retained IC temporal components were low-pass filtered at 2 Hz for stimulation data and 0.1 Hz for resting-state data, Z-scored, and min–max normalized. The target number of ICs was 30. For the somatosensory and resting-state datasets, ICs were removed iteratively until the maximum variance inflation factor (VIF) fell below 10. ROI-based features were extracted from 26 cortical regions of the standard atlas and processed identically for comparison (Supplementary Methods).

### Dataset construction and selection

Each sample paired the time series of all features over a 1.5 s window (31 frames for sensory stimulation data or 46 frames for resting-state data) with the binary pupil label at the final frame (the pupil-labeled frame). Sessions in which the pupil reached the threshold before the pre-peak window or never reached it after stimulation were excluded (Supplementary Methods); the pre-peak window for each condition is given in the Results.

### Decoder construction and evaluation

Three architectures were evaluated: Gated Recurrent Unit (GRU), Long Short-Term Memory (LSTM), and Linear Support Vector Machine (Linear SVM). The GRU models followed a previous study (Ajioka et al. 2024), with a single GRU layer (32 units), a fully connected layer (1 unit), and a sigmoid output layer (Supplementary Methods). Each model was trained for 30 epochs (somatosensory stimulation) or 10 epochs (other conditions), and the model with the lowest validation loss was selected.

The same three mice were used in all three sensory stimulation experiments. Data from each mouse were randomly divided into five equal-sized partitions and pooled across animals to form five balanced folds, with partitioning at the session level such that all samples from a session were assigned to the same fold. Five-fold cross-validation yielded five trained models per architecture; null models were trained identically, except that labels were randomly shuffled within sessions. The area under the ROC curve (ROC-AUC) was the primary metric of discriminability, and for visualization decoder output was binarized at the threshold maximizing the Youden index.

### Deep SHAP and back-mapping

Feature contributions to GRU models were quantified using Deep SHAP (Lundberg and Lee 2017): SHAP values were calculated for the first sample in each validation trial that yielded a positive prediction, using every 20th sample of the entire validation dataset as the background reference distribution and averaged across models. SHAP values for each IC feature were back-mapped onto cortical space as weights for a linear combination of the corresponding ICA spatial components. Mapped SHAP arrays from the three sensory datasets were normalized by the sum of their absolute values and jointly subjected to PCA, yielding 10 components (Supplementary Fig. 12A), whose spatial components were used to calculate PCA temporal components for null models and resting-state data.

### Statistical analyses

Statistical analyses were performed in Python using the SciPy library. Comparisons used paired Student’s t-tests, Wilcoxon signed-rank tests, or Mann–Whitney U tests, as specified in the figure legends; nonparametric tests were used when normality could not be assumed, and formal normality tests were not performed (Supplementary Methods). Where the legends indicate it, P-values were Holm-adjusted within the pairwise comparisons of each panel; all other comparisons were unadjusted. N denotes the number of mice and n the number of trials or recording sessions, except for machine-learning model evaluation, where n is the number of trained models (n = 5). Differences were considered statistically significant at P < 0.05. Analysis code was written in-house (see Code Availability) and executed in Python 3.6 (decoders) or Python 3.11.

## Results

### Simultaneous monitoring of pupil dilation and cortical activity

Pupil dilation and cortical activity during somatosensory stimulation (hindpaw electrical stimulation) were monitored simultaneously by imaging the eye (Fig. 1A; (−), before and Stim, during stimulation) and by two-wavelength wide-field Ca²⁺ imaging using GCaMP6f (Fig. 1B). Average traces of pupil diameter and Ca²⁺-dependent components in the primary somatosensory cortex (SSp) showed an increase in cortical activity followed by pupil dilation (Makino et al. 2017) (Fig. 1C). Both averages were more prominent for the 42 sessions (35.3%) selected for decoder training (Fig. 1C, right, Select) than for all sessions (Fig. 1C, left, All). Frequency components increased in the 0.5–1.5 Hz band after stimulation, more prominently after 3 s in Select (Fig. 1D), and cortical activity correlated positively with pupil diameter immediately after stimulation and negatively thereafter, again more prominently in Select (Fig. 1E).

**Figure 1.**
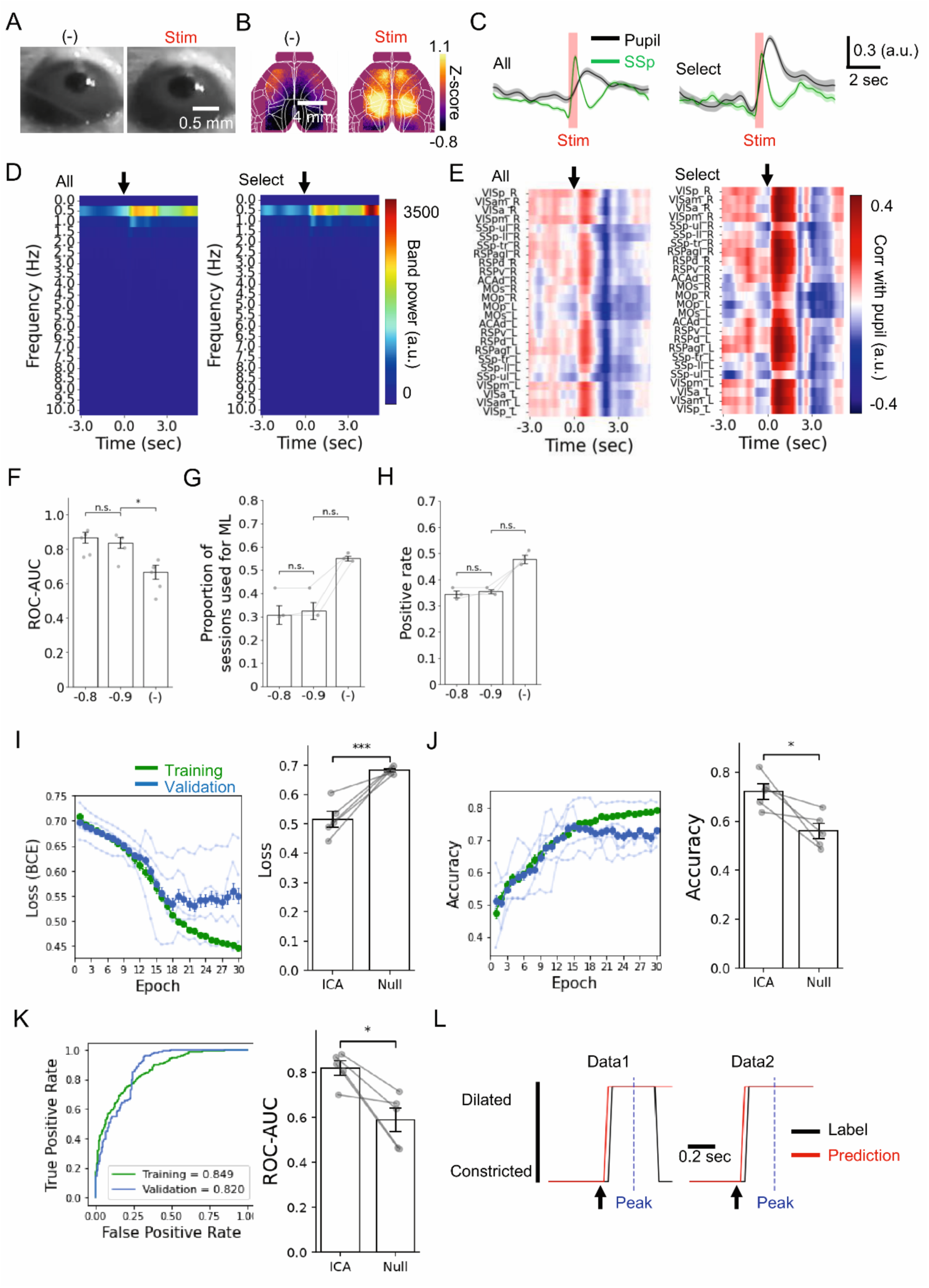
RNN-based decoder predicts somatosensory-evoked pupil dilation from cortical activity. *A*, Representative pupil images before (−) and during somatosensory stimulation, Stim. ***B*,** Representative GCaMP6f fluorescence images after hemodynamic correction. ***C,*** Time course of pupil diameter (Pupil) and primary somatosensory cortex activity (SSp). Mean ± SD of all datasets, all (N = 3 mice, n = 119 sessions), and those selected for machine learning, Select (N = 3 mice, n = 42 sessions). ***D,*** Frequency components of cortical activity in response to the stimuli (arrows). Bin size, 0.05 s. ***E,*** Pearson correlations between cortical activity and pupil diameter across cortical regions. ***F,*** Effect of input time range on model performance assessed by the area under the ROC curve (ROC-AUC). Time ranges began 0.8 s (−0.8), 0.9 s (−0.9) or no constraint (−) before the pupil diameter peak, which was detected 0.5 s or later after stimulus onset. Median ± SE (n = 5 models). Mann–Whitney U test with Holm correction; n.s. P > 0.05, *P < 0.05. ***G*** Effect of input time range on the proportion of sessions selected for machine learning. ***H,*** Effect of input time range on the positive label rate (Positive rate). Mean ± SE (N = 3 mice) for G and H. Wilcoxon signed-rank test with Holm correction; n.s. P > 0.05. ***I,*** Prediction by RNN-based decoder assessed by loss (BCE; binary cross-entropy). Loss changes in training (training) and validation (validation) datasets over 30 epochs (epoch) (*left*) and comparison with a null model trained on within-session target-shuffled datasets (*right*). The dataset was randomly divided into five equal folds: four were used for training and one for validation using a leave-one-out approach, generating models (n = 5). ***J,*** Prediction assessed by accuracy. ***K,*** Prediction assessed by ROC curves. ROC curves from a representative decoder and the mean ROC-AUC of the training and validation datasets across the five models (*left*). Comparison of ROC-AUC with the null model (*right*). Mean ± SE (n = 5 models). Paired t-test; *P < 0.05, ***P < 0.005 for I, J, and K. ***L,*** Observed (label) and predicted (Prediction) pupil states in two representative sessions (Data1, Data2). Observed states were classified using a threshold of 0.6 applied to the normalized pupil diameter to define the dilated state for machine learning labels. Predicted states were classified using a threshold of 0.431, which maximized the difference between the true positive rate and false positive rate on the ROC curve. Alt text: Panels A and B show pupil images and cortical calcium activity maps before and during somatosensory stimulation. C shows the cortical activity preceding pupil dilation. D and E show stimulus-related changes in cortical frequency power and region-specific correlations with the pupil diameter. F through H show that restricting input time windows improves decoder performance, but reduces the proportion of selected sessions and positive labels. I through K show lower loss, higher accuracy, and higher ROC-AUC than the shuffled-label null models. L compared the predicted and observed pupil states in two representative sessions.

### Independent component analysis of cortical activity

Decoder analysis requires statistically independent features, so ROI-and ICA-based features were compared using the variance inflation factor (VIF) (Dormann et al. 2013; Aas et al. 2021; Molnar et al. 2022). For the somatosensory dataset, the maximum VIF before stimulation onset was 7.8 for 10 independent components (ICs), but was significantly larger and exceeded 10 for 12 and 14 ICs, indicative of severe multicollinearity (O’brien 2007; Dormann et al. 2013), and reached 1954.16 for ROI-based features (Supplementary Figs. 2 and 3A–C). The decoder was trained using 10 ICs.

### An RNN decoder predicts somatosensory-evoked pupil dilation from cortical activity

A gated recurrent unit (GRU), a variant of RNN (Elman 1990; Cho et al. 2014), was employed as the decoder architecture, and cortical data from stimulation onset to 0.5 s after the pupil diameter peak were included. To minimize the influence of spontaneous dilation driven by tonic locus coeruleus (LC) activation, sessions exhibiting dilation within a pre-peak window of 0.8 s (−0.8) or 0.9 s (−0.9) were excluded; both constraints significantly raised ROC-AUC relative to no constraint (−) and did not differ from each other, while reducing the proportion of sessions selected (55.4% versus 35.3% and 34.4%) and of dilation events within them (48.2% versus 35.7% and 34.7%) (Fig. 1F–H). The temporal range was therefore set from −0.9 to +0.5 s relative to the pupil peak.

Validation loss and accuracy saturated between epochs 15 and 18 (Fig. 1I–J, left), and a null model with labels shuffled within each session showed significantly higher loss and lower accuracy (Fig. 1I–J, right). ROC-AUC exceeded 0.8 for both training and validation data and was significantly higher than for the null decoder (Fig. 1K). In two representative sessions, the dilation label (Label) closely matched the model’s classification at the ROC-optimal threshold (Prediction) (Fig. 1L). Therefore, the GRU decoder predicts stimulation-evoked pupil dilation, suggesting that specific spatiotemporal cortical patterns precede it.

### An RNN decoder predicts visual pattern-evoked pupil dilation

Visual stimulation employed drifting gratings, which are constant in light intensity, elicit minimal light-induced pupil responses, and are commonly used to assess visual cognition, such as orientation, spatial frequency selectivity, and contrast sensitivity (Prusky et al. 2004; Niell and Stryker 2008). The two orientation change events (ex1 and ex2) were used as stimuli (Fig. 2A). This stimulation elevated Ca²⁺-dependent components of the primary visual cortex (VISp) (Fig. 2B). All sessions (n = 741) showed pupil dilation and increased VISp activity (Fig. 2C, left), while 143 sessions (19.3%) selected for decoder training showed more pronounced dilation (Fig. 2C, right). Spectral power increased in the 0.5–1.5 Hz band in both groups (Fig. 2D). All sessions showed a positive correlation between cortical activity and pupil diameter from approximately 1 s after stimulation, whereas selected sessions showed correlations that were positive before and immediately after onset, then negative, then positive again (Fig. 2E); the pre-stimulus positive correlation may indicate that ongoing activity modulated pupil responsiveness. ICA yielded 23 independent components (Supplementary Fig. 4) with low multicollinearity (maximum VIF = 6.5; Supplementary Fig. 3D). ROC-AUC did not differ significantly between ROI-and ICA-based features or among GRU, LSTM, and linear SVM, although GRU and LSTM tended to exceed linear SVM (Fig. 2F). A pre-peak cutoff of −1.5 s tended to yield higher ROC-AUC while reducing the number of selected sessions and the proportion of positive labels (Fig. 2G–I), and a pupil label threshold of 0.6 was optimal (Supplementary Fig. 5). A GRU decoder was therefore trained on the dataset spanning −1.5 to +0.5 s. Validation loss decreased by epoch 4 and was significantly lower than that of the null model (P = 0.0216; Fig. 2J), although accuracy did not differ (P = 0.9737; Fig. 2K). Validation ROC-AUC exceeded 0.6, which was significantly higher than that of the null model (P = 0.0048; Fig. 2L), and prediction capability was confirmed in two representative sessions (Fig. 2M), indicating that visual pattern-evoked dilation is preceded by specific cortical activity patterns.

**Figure 2.**
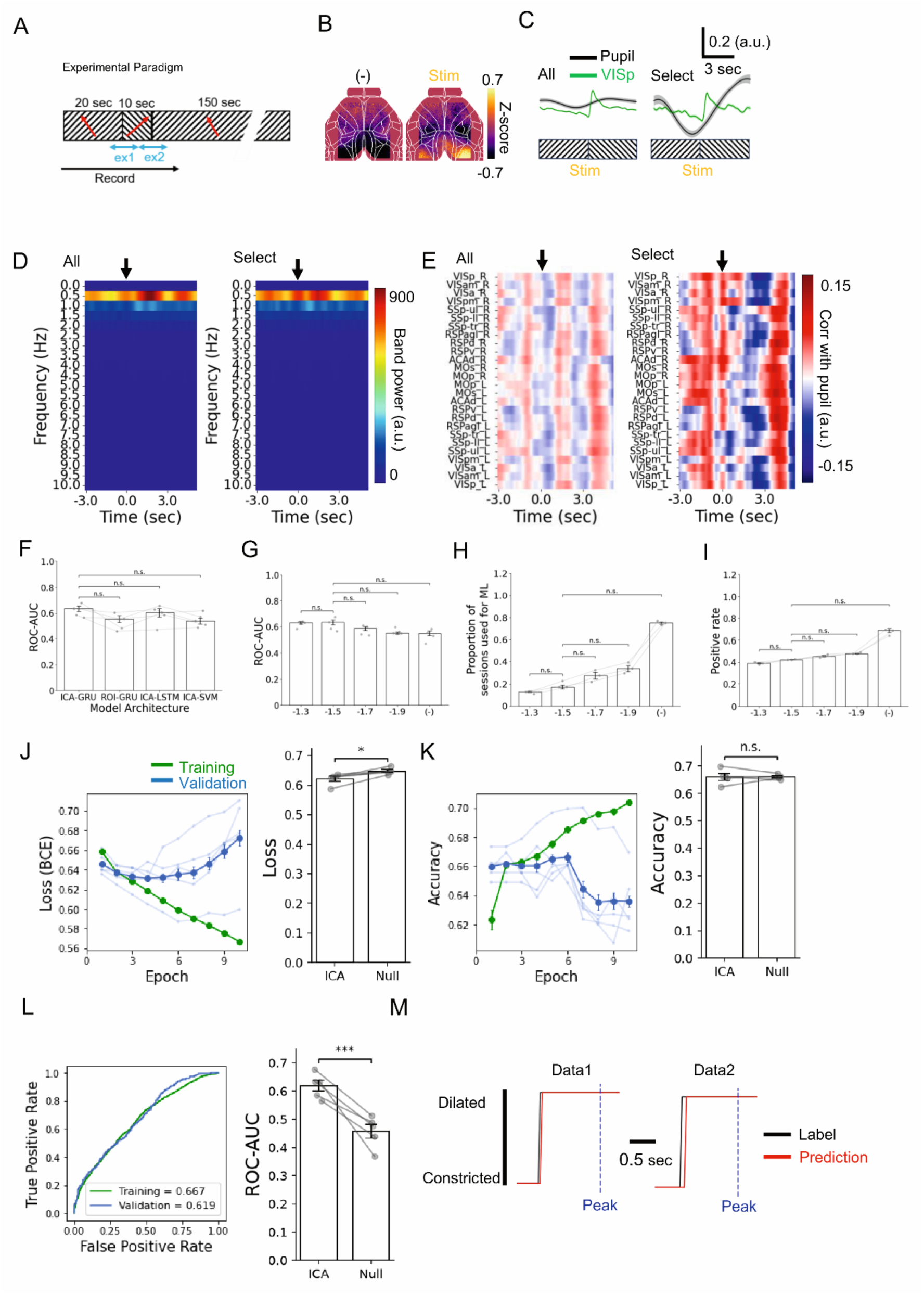
RNN-based decoder predicts pupil dilation evoked by visual pattern stimulation from cortical activity. ***A,*** Schematic of visual pattern stimulation. The mice were head-fixed on a treadmill with a monitor displaying a grating pattern (Lateral View). A mirror reflecting the monitor image and an infrared camera for pupil imaging (Camera1) were positioned in front of the animal (Top View). The direction of the grating flow was changed over 10 s, and pupil and cortical activity were recorded for 40 s starting 20 s prior to the change (Experimental Paradigm). Both grating direction change events (ex1 and ex2) are included in the dataset. ***B,*** Representative cortical activity before (−) and during stimulation, Stim. ***C,*** Time course of pupil diameter (Pupil) and primary visual cortex activity (VISp). *Left:* Mean ± SD of all datasets (all; N = 3 mice, n = 741 sessions). *Right:* those selected for machine learning (Select; N = 3 mice, n = 143 sessions). ***D,*** Frequency components of cortical activity in response to the stimuli (arrows). Bin size, 0.05 s. ***E,*** Pearson correlations between cortical activity and pupil diameter across cortical regions. ***F,*** Effect of training conditions assessed by the area under the ROC curve (ROC-AUC). The input datasets (ICA vs. ROI) and decoder architectures (GRU, LSTM, and linear SVM) were compared. Median ± SE (n = 5 models). Wilcoxon signed-rank test with Holm correction; n.s., P > 0.05. ***G,*** Effect of input time range on model performance assessed by ROC-AUC. Median ± SE (n = 5 models). Mann–Whitney U test with Holm correction; n.s., P > 0.05. Time ranges began 1.3 s (−1.3), 1.5 s (−1.5), 1.7 s (−1.7), 1.9 s (−1.9) or no constraint (−) before the pupil diameter peak, which was detected 0.5 s or later after stimulus onset. ***H***, Effect of input time range on the proportion of sessions selected for machine learning. ***I,*** Effect of input time range on positive label rate (Positive rate). Mean ± SE (N = 3 mice) for H and I. Wilcoxon signed-rank test with Holm correction; n.s. P > 0.05. ***J,*** Prediction by RNN-based decoder assessed by loss (BCE; binary cross-entropy). Loss changes in training and validation datasets over 10 epochs (*left*) and comparison with a null model trained on within-session target-shuffled datasets (*right*). The dataset was randomly divided into five equal folds: four were used for training and one for validation using a leave-one-out approach, generating models (n = 5). ***K,*** Prediction assessed by accuracy. ***L,*** Prediction assessed by ROC curves. ROC curves from a representative decoder and the mean ROC-AUC of the training and validation datasets across the five models (*left*). Comparison of ROC-AUC with the null model (*right*). Mean ± SE (n = 5 models). Paired t-test; n.s. P > 0.05, *P < 0.05, ***P < 0.005 for J, K and L. ***M,*** Observed (label) and predicted (Prediction) pupil states in two representative sessions (Data1, Data2). Observed states were classified using a threshold of 0.6 applied to the normalized pupil diameter to define the dilated state for machine learning labels. Predicted states were classified using a threshold of 0.668, which maximized the difference between the true positive rate and false positive rate on the ROC curve. Alt text: Multi-panel figure showing visual pattern stimulation, cortical activity maps, and pupil responses. Panels D and E show the stimulus-related frequency power and regional correlations with pupil diameter. F and G compare the decoder performance across model architectures and input time windows, showing no significant differences. H and I show increased session inclusion and positive label rates with unrestricted input windows. J through L compare RNN-based decoding with shuffled-label null models, showing lower loss and higher ROC-AUC, but similar accuracy. M shows the predicted pupil-state transitions closely matching the observed transitions in two representative sessions.

### An RNN decoder predicts visual context-evoked pupil dilation

For visual context stimulation, an oddball paradigm using the same orientation change was adopted, with control stimuli of random orientations, followed by test stimuli comprising redundant stimuli and a single deviant stimulus (Fig. 3A). Deviant stimuli preferentially elevated cortical Ca²⁺-dependent components relative to redundant stimuli (Fig. 3B). All sessions (n = 1107) showed no prominent pupil dilation or VISp activity (Fig. 3C, left), and neither the magnitude and velocity of pupil dilation nor MOs activity differed significantly from control responses (Supplementary Fig. 6A–C). However, the 79 sessions (7.1%) selected for decoder training showed dilation that began during the redundant stimulus and became more pronounced during the deviant stimulus (Fig. 3C, right). Spectral power increased in the 0.5–2.0 Hz band in both groups (Fig. 3D); across all sessions the correlation between cortical activity and pupil diameter became moderately negative after deviant stimulation, whereas in selected sessions correlations of primary and higher-order visual cortex activity were more pronounced from 0.5 s after onset (Fig. 3E). ICA yielded 25 independent components (Supplementary Fig. 7) with low multicollinearity (maximum VIF = 7.7; Supplementary Fig. 3E). Since a pre-peak cutoff of −1.2 s tended to improve model performance despite reducing the number of selected sessions and the proportion of positive labels (Fig. 3F–H), a GRU decoder was trained on the dataset spanning −1.2 to +0.5 s. Validation loss decreased by epoch 4 and was significantly lower than that of the null model (P = 0.0117; Fig. 3I), accuracy was significantly higher (P = 0.0122; Fig. 3J), and ROC-AUC exceeded 0.6 and was significantly higher (P = 0.0044; Fig. 3K). Prediction capability was confirmed in two representative sessions (Fig. 3L), indicating that pupil dilation evoked by visual context stimulation is also preceded by specific cortical activity patterns.

**Figure 3.**
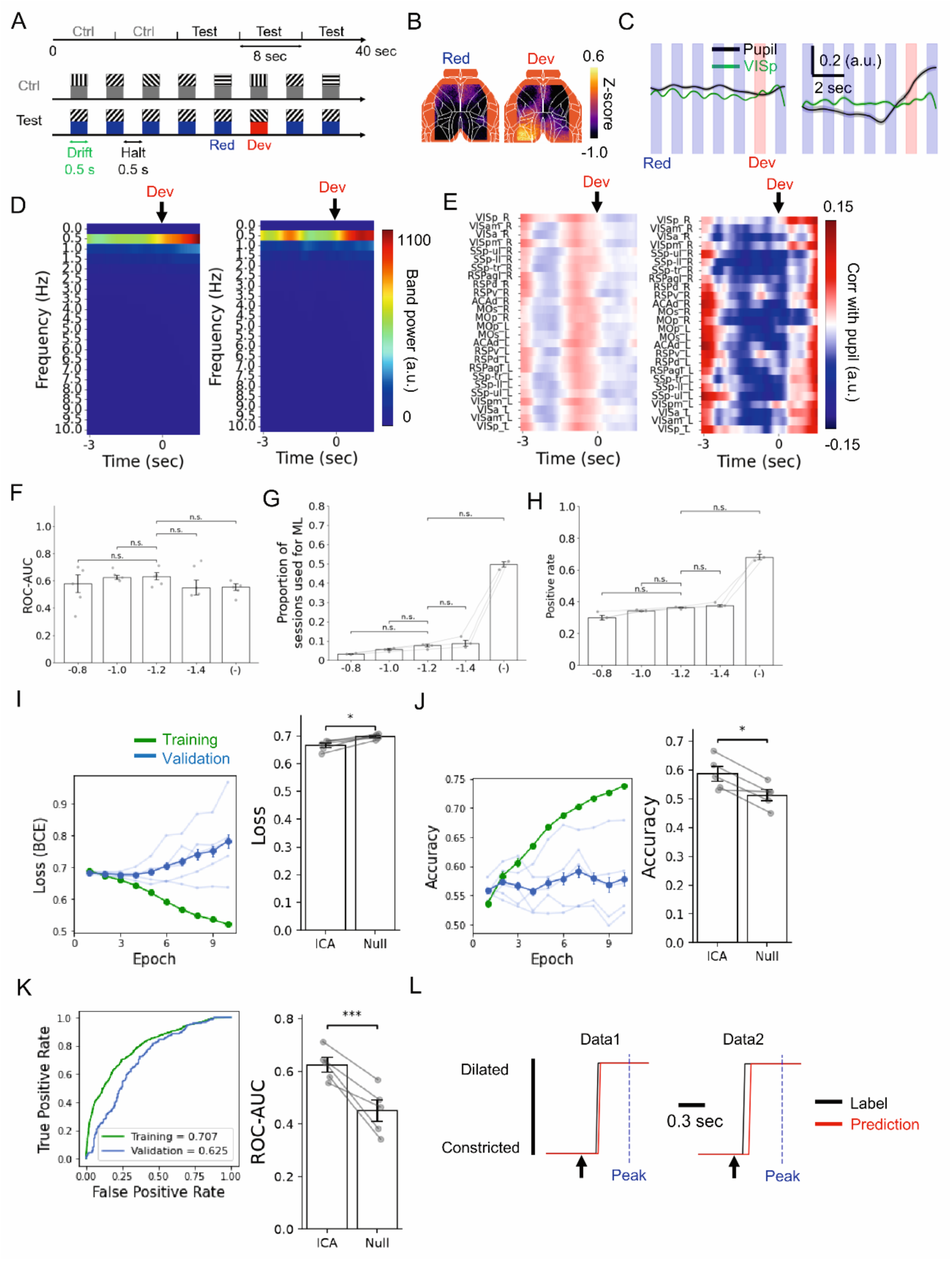
RNN-based decoder predicts pupil dilation evoked by visual context stimulation from cortical activity. ***A,*** Schematic of the visual oddball task for context stimulation. Each block consisted of a 0.5 s halt followed by a 0.5 s drift. Mice were subjected to two repetitions of control stimulation (Ctrl), consisting of eight blocks with random drift directions, followed by three repetitions of a test cycle, comprising five blocks with redundant drift directions (Red), one block with deviant drift directions (Dev), and two blocks with redundant drift directions. ***B,*** Representative cortical activity during redundant (red) and deviant (Dev) stimulation. ***C,*** Time course of pupil diameter (Pupil) and primary visual cortex activity (VISp). *Left:* Mean ± SD of all datasets (all; N = 3 mice, n = 1107 sessions). *Right:* those selected for machine learning (Select; N = 3 mice, n = 79 sessions). ***D,*** Frequency components of the cortical activity in response to deviant stimuli (arrows). Bin size, 0.05 s. ***E,*** Pearson correlations between cortical activity and pupil diameter across cortical regions. ***F,*** Effect of input time range on model performance assessed by the area under the ROC curve (ROC-AUC). Time ranges began 0.8 s (−0.8), 1.0 s (−1.0), 1.2 s (−1.2), 1.4 s (−1.4) or no constraint (−) before the pupil diameter peak, which was detected 0.5 s or later after stimulus onset. Median ± SE (n = 5 models). Mann– Whitney U test with Holm correction; n.s., P > 0.05. ***G*** Effect of input time range on the proportion of sessions selected for machine learning. ***H,*** Effect of input time range on the positive label rate (Positive rate). Mean ± SE (N = 3 mice) for G and H. Wilcoxon signed-rank test with Holm correction; n.s. P > 0.05. ***I,*** Prediction by RNN-based decoder assessed by loss (BCE; binary cross-entropy). Loss changes in training (training) and validation (validation) datasets over 10 epochs (epoch) (*left*) and comparison with a null model trained on within-session target-shuffled datasets (*right*). The dataset was randomly divided into five equal folds: four were used for training and one for validation using a leave-one-out approach, generating models (n = 5). ***J,*** Prediction assessed by accuracy. ***K,*** Prediction assessed by ROC curves. ROC curves from a representative decoder and the mean ROC-AUC of the training and validation datasets across the five models (*left*). Comparison of ROC-AUC with the null model (*right*). Mean ± SE (n = 5 models). Paired t-test; *P < 0.05, ***P < 0.005 for I, J, and K. ***L,*** Observed (label) and predicted (Prediction) pupil states in two representative sessions (Data1, Data2). Observed states were classified using a threshold of 0.6 applied to the normalized pupil diameter to define the dilated state for machine learning labels. Predicted states were classified using a threshold of 0.431, which maximized the difference between the true positive rate and false positive rate on the ROC curve. Alt text: Twelve-panel figure showing the visual oddball task and cortical decoding of pupil dilation. A and B show the stimulus sequences and cortical activity maps for redundant and deviant stimuli, respectively. C shows pupil dilation following deviant stimulation, along with visual cortical activity. D and E show the frequency power and regional correlations with pupil diameter. F through H show similar ROC-AUC across the input windows, whereas unrestricted windows include more sessions and positive labels. I through K show lower loss, higher accuracy, and higher ROC-AUC than the shuffled-label null models. L shows closely aligned observed and predicted pupil state transitions.

### Feature importance implicates cortical activity preceding pupil dilation

Because permutation importance quantifies only the magnitude of a feature’s contribution and not its direction, contributions were analyzed using Deep SHAP (Lundberg and Lee 2017), which resolves the direction of each IC’s contribution in each time window. A model of spontaneous pupil dilation during rest was constructed from an existing dataset (Kondo et al. 2025) for comparison. ICA yielded 10 independent components with low multicollinearity (maximum VIF = 9.0), and the decoder, trained and evaluated as for the sensory conditions, predicted spontaneous dilation with ROC-AUC significantly above the null model when the data were low-pass filtered at 0.1 Hz; neither the number of input ICs nor the start time of the input window affected performance (Supplementary Figs. 3F–G and 8).

Holm correction was applied throughout because SHAP values capture inter-regional relationships. In the somatosensory model, IC5, spanning the retrosplenial cortex (RSP) to higher-order visual areas, showed elevated SHAP values from −0.5 to 0.0 s (P = 0.013, 0.018 and 0.012 for −0.5 to −0.3, −0.3 to −0.1 and −0.1 to 0.0 s) (Fig. 4A–B Som; Supplementary Fig. 9A). In the visual pattern model, values were elevated for IC1 (motor cortex) at −0.3 to −0.1 s (P = 0.0002) and for IC5 (anterior cingulate area, dorsal part; ACAd) and IC15 (ACAd and RSP) at −0.1 to 0.0 s (P = 0.002 and 0.0015) (Fig. 4A–B VP; Supplementary Fig. 9B). In the visual context model, IC4 (visual cortex and RSP) contributed strongly at −0.3 to −0.1 s (P = 0.0008) and −0.1 to 0.0 s (P = 0.0030) (Fig. 4A–B VC; Supplementary Fig. 9C). In the spontaneous model, IC7 (ACAd) increased at −1.0 to −0.5 s (P = 0.0208) and −0.5 to −0.3 s (P = 0.0393), followed at −0.3 to −0.1 s by IC4 (MOs and SSp; P = 0.0337) and IC6 (RSP; P = 0.0395) (Fig. 4A–B Spont; Supplementary Fig. 9D). Decoders trained on high-|SHAP| IC subsets (top 5, 10, and 13 ICs for somatosensory, visual pattern, and visual context stimulation) yielded ROC-AUC values 43.1% (P = 0.0005), 19.1% (P = 0.0181), and 37.1% (P = 0.0233) higher than those trained on the corresponding low-|SHAP| subsets (Supplementary Fig. 10A– B, D–E, G–H). For somatosensory stimulation, features within the −1.50 to −0.25 s window yielded an ROC-AUC 23.6% higher than null features (P = 0.0212; Supplementary Fig. 10C), whereas for both visual conditions models excluding features at 0.0 s did not differ from their null-feature models (Supplementary Fig. 10F and I).

**Figure 4.**
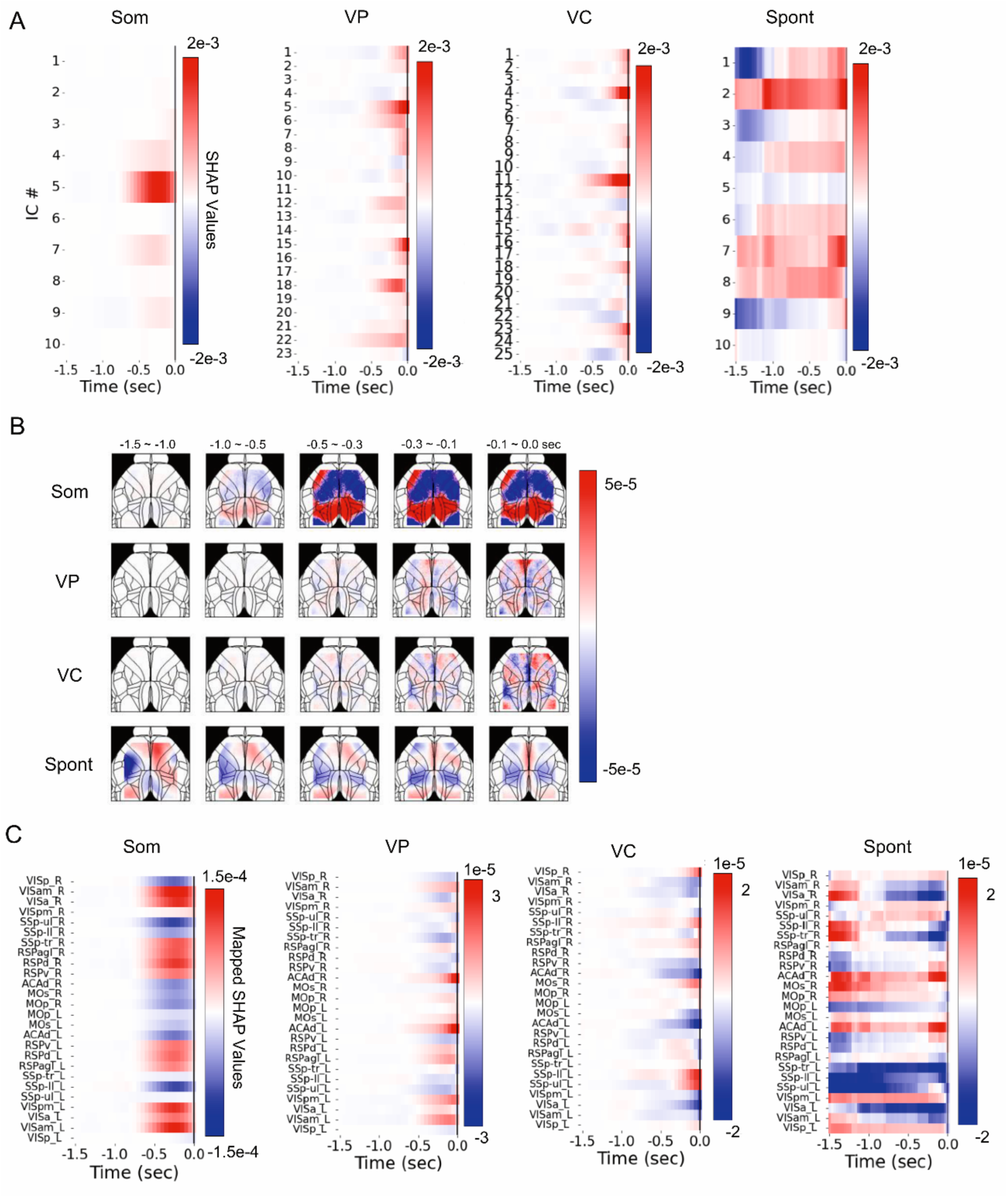
SHAP analysis identifies contributions of the anterior cingulate cortex, secondary motor area, and retrosplenial cortex in evoked and spontaneous pupil dilation. ***A,*** Heat map of time-dependent change in the SHAP value of independent components (ICs). The ICs were extracted from the RNN decoder using validation data. Time is shown relative to the pupil-labeled frame (0.0 s), that is, the frame carrying the binary pupil label predicted by the decoder. Somatosensory stimulation (Som), visual pattern stimulation (VP), visual context stimulation (VC), and spontaneous pupil dilation (Spont, N=25 mice, n=923 sessions, 5 models). ***B,*** Spatial distribution of SHAP values across time windows. The SHAP values were mapped onto a standard brain atlas using ICA spatial components. The time windows are shown relative to the pupil-labeled frame (0.0 s). ***C,*** Heat maps of time-dependent changes in mapped SHAP values across brain regions. Alt text: Three-panel figure showing temporal and spatial SHAP contributions to pupil dilation prediction across somatosensory, visual pattern, visual context, and spontaneous conditions. A shows the time-dependent SHAP values across the independent components, with distinct patterns among the conditions. B maps the positive and negative SHAP contributions across cortical regions and time windows preceding the predicted pupil state. C shows region-specific temporal SHAP patterns with prominent contributions from the retrosplenial, anterior cingulate, and motor cortical regions. Somatosensory stimulation shows pronounced contributions near the pupil-labeled frame, whereas spontaneous pupil dilation exhibits more temporally distributed patterns.

SHAP values were then mapped onto each pixel using ICA spatial components so that high mapped values indicate a shift of the model output toward the dilation class. The somatosensory model showed higher mapped SHAP values than the null model across a broad region encompassing RSP and higher-order visual areas during the three windows from −0.5 to 0.0 s (Fig. 4C Som; Supplementary Fig. 11A). The visual pattern model showed elevated values in RSP and the anterior visual area (VISa) at −0.3 to −0.1 s, and in RSP, the posteromedial visual area (VISpm), ACAd and the secondary motor cortex (MOs) at −0.1 to 0.0 s (Fig. 4C VP; Supplementary Fig. 11B); the visual context model showed elevated values specifically in MOs at −0.1 to 0.0 s (Fig. 4C VC; Supplementary Fig. 11C), and the spontaneous model showed negative values in the primary somatosensory area-trunk (SSp-tr) and VISa at −0.3 to −0.1 and −0.1 to 0.0 s (Fig. 4C Spont; Supplementary Fig. 11D).

### Cortical dynamics associated with evoked and spontaneous pupil dilation

To visualize the cortical dynamics associated with pupil dilation, PCA was applied to the mapped SHAP values, normalized by the sum of their absolute values. The top three components accounted for 53.1% of the variance (Supplementary Fig. 12A) and captured distinct spatial patterns: PC1 spanned V1, the somatosensory cortex, and anterior MOs; PC2 was centered on ACAd and the RSP–higher-order visual area region; and PC3 spanned RSP, higher-order visual areas, and posterior MOs (Fig. 5A). The 1.5 s trajectories in PC1–PC3 space revealed stimulus-specific dynamics (Fig. 5B). PC2 and PC3 increased during −0.5 to −0.3 s for somatosensory stimulation (Fig. 5C, Som; Supplementary Fig. 12B), whereas PC1 increased during −1.0 to −0.5 s and −0.5 to −0.3 s for visual pattern stimulation (Fig. 5C, VP; Supplementary Fig. 12C). Visual context stimulation showed the same PC1 increase plus PC2 and PC3 increases during −0.3 to 0.0 s, with PC1 tending to remain elevated from −0.5 s onward, although high inter-trial variability made definitive evaluation difficult (Fig. 5C, VC; Supplementary Fig. 12D). For spontaneous dilation, PC3 decreased progressively from −1.0 to −0.3 s (Fig. 5C, Spont; Supplementary Fig. 12E). When temporal components extracted using the PCA spatial components were used as decoder input, pupil state was significantly predicted for all three stimulation conditions but not for spontaneous dilation (Fig. 5D).

**Figure 5.**
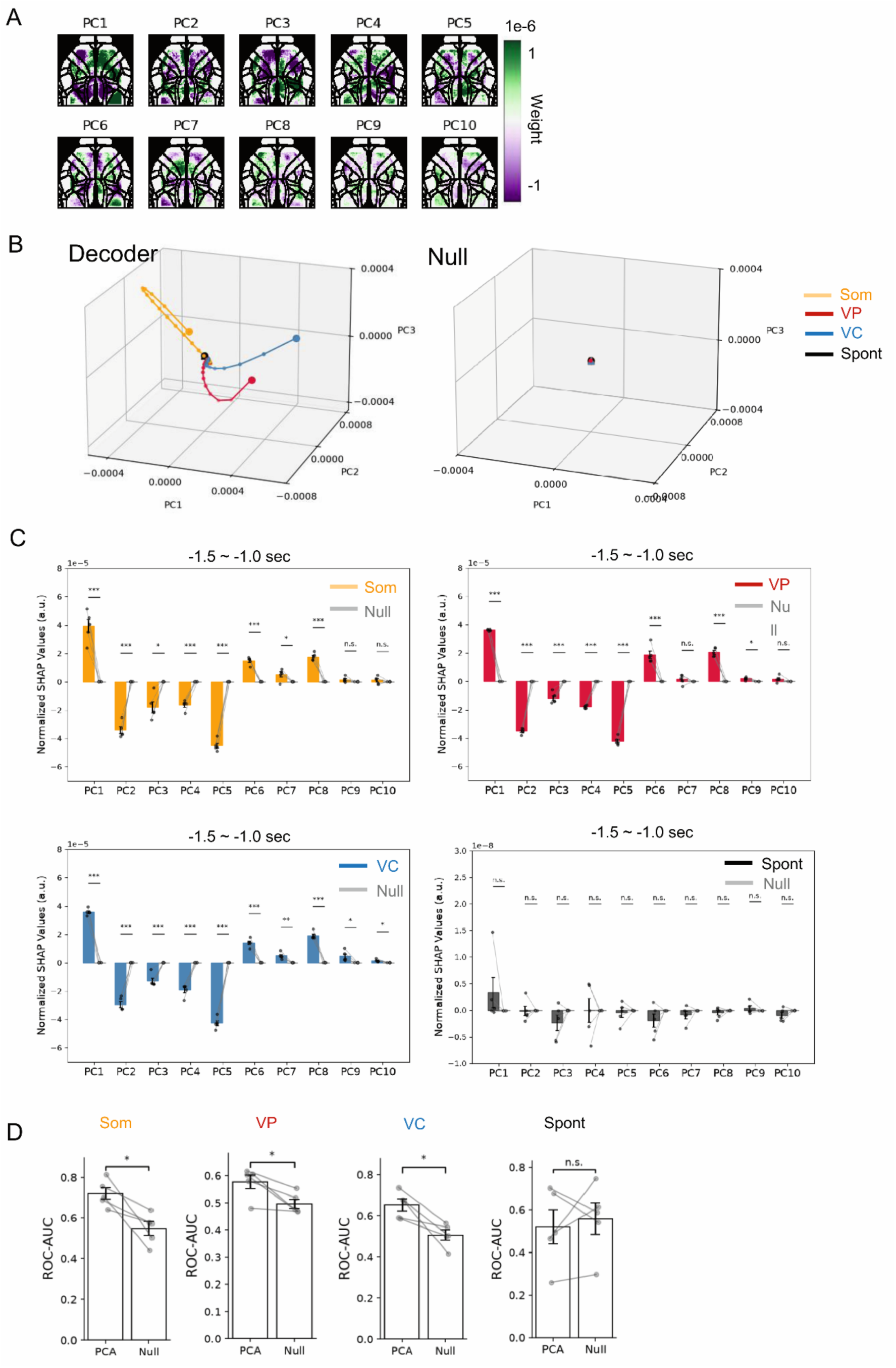
Cortical activity underlying sensory-evoked pupil dilation consists of shared and stimulus-specific components. ***A,*** PCA spatial components (PC1 – PC10) derived from the PCA of SHAP values projected onto a standard brain atlas. Projected SHAP values were obtained from five models trained on somatosensory, visual pattern, or visual context stimulation data as inputs and normalized by the sum of absolute values before PCA. Color indicates the component weight. ***B,*** Trajectories of the top three PCA temporal components within the −1.5 to 0.0 s time window relative to the pupil-labeled frame (0.0 s) for decoder (*left*) and null (*right*) models. The start and end points are indicated by the triangles and circles, respectively. Temporal resolution: 0.05 s The Som, VP, and VC were 0.03 s for Spont. Consecutive timeframes were connected using straight lines. ***C,*** Normalized SHAP values of PCA temporal components in the indicated time window compared with the null model. ***D,*** ROC-AUC of ROC curves for RNN decoders with temporal components extracted from hemodynamic-corrected images based on PCA spatial components as input features. Mean ± standard error of n = 5 models. Paired t-test (uncorrected). n.s. P > 0.05, *P < 0.05, **P < 0.01, ***P < 0.005. Alt text: Four-panel figure showing PCA of cortical SHAP contributions across sensory-evoked and spontaneous pupil dilation. A maps the spatial weights of the ten principal components onto the cortex. B shows distinct temporal trajectories for somatosensory, visual pattern, and visual context conditions, while null-model trajectories are clustered near the origin. C shows significant positive and negative SHAP contributions relative to null models during sensory-evoked, but not spontaneous pupil dilation. D shows significantly higher ROC-AUC than the null models for all three sensory conditions, but not for spontaneous pupil dilation.

## Discussion

We tested whether sensory-evoked pupil dilation is preceded by specific cortical activity involved in sensory perception, and whether spontaneous dilation is preceded by a different pattern. RNN decoders predicted dilation from ICA-derived cortical components, and SHAP-based feature importance analysis identified the most predictive regions across three sensory conditions and an open-source resting-state dataset; our imaging parameters are broadly consistent with previous wide-field studies (Musall et al. 2019; Couto et al. 2021; Musall et al. 2023; Raut et al. 2025).

### Cortical signatures of sensory-evoked pupil dilation

Somatosensory stimulation evoked a widespread increase in cortical [Ca²⁺]ᵢ coincident with pupil dilation, consistent with previous reports (Slezak et al. 2019; Lee et al. 2020), and SHAP values were distributed across the retrosplenial cortex (RSP) and higher-order visual areas, supporting spatial navigation, memory retrieval, and context representation (Vann et al. 2009; Murakami et al. 2015) and directional and visuospatial processing (Marshel et al. 2011; Andermann et al. 2011), respectively. Hindpaw stimulation elicits locus coeruleus (LC) spiking preceding somatosensory cortex depolarization and pupil dilation (Sara and Bouret 2012; Yang et al. 2021), so our findings may partly reflect this LC-driven response. In contrast, whisker deflection reduces inter-areal functional connectivity (Mitelut et al. 2022), possibly because electrical stimulation also activates the nociceptive system (Grivet et al. 2025). Non-nociceptive stimuli may suit future studies.

Under visual pattern stimulation, mapped SHAP increases in RSP and the anterior visual area (VISa) preceded those in the dorsal anterior cingulate cortex (ACAd) and motor cortex. This matches reported [Ca²⁺]ᵢ elevations in RSP, MOs, and ACA in response to drifting gratings (Murakami et al. 2015) and supports a model in which RSP, as part of the dorsal visual stream, is driven by higher-order visual input, with MOs and ACA as downstream targets (Wang et al. 2012); ACA in turn sends feedback projections that modulate visual discrimination (Zhang et al. 2014; Zingg et al. 2014). Their localization within visual-related areas suggests that the activity contributing to dilation reflects visual perception upstream of, rather than downstream from, LC activation.

Pupil dilation to novel visual context stimuli was not significant at the group level; however, models trained on sessions exhibiting dilation predicted above chance, with MOs showing high mapped SHAP values, although MOs itself did not respond significantly. A previous study using the same paradigm reported no significant dilation and showed that context selectivity was diminished by suppressing prefrontal input (Hamm et al. 2021; Homann et al. 2022). The prominence of MOs raises the possibility that context changes were processed as prediction errors, although orientation changes delivered once per 8 s may also have induced context-independent responses. Retrosplenial involvement is thus common to somatosensory and visual stimulation, whereas prominent MOs involvement is restricted to visual conditions.

### Spontaneous pupil dilation

Spontaneous dilation could not be decoded above chance from activity low-pass filtered at 2 Hz; a 0.1 Hz filter was required. This supports the proposal that LC activity underlying dilation is transient during sensory stimulation but sustained during spontaneous arousal fluctuations (Aston-Jones and Cohen 2005), and that coupling between spontaneous dilation and cortical activity is restricted to frequencies below 1 Hz (Turner et al. 2023; Raut et al. 2025). SHAP increases in the medial anterior cortex (IC7) preceded those in the medial posterior region (IC6) and MOs (IC4), which, to our knowledge, has not been previously reported. Earlier work removed arousal-related co-variation between pupil size and brain activity by linear regression (Salkoff et al. 2020; West et al. 2024). More recently, cortical activity was predicted more accurately from a low-dimensional arousal state space, indicating that pupil–cortex coupling is nonlinear (Raut et al. 2025). Nevertheless, wide-field imaging is restricted to superficial layers and does not capture salience network structures such as the insular cortex or claustrum.

### Cortical dynamics and the conditions for successful decoding

PCA of mapped SHAP values dissociated somatosensory from visual conditions along PC1, and PC2 was prominent in both somatosensory and visual pattern stimulations. The prolonged elevation of PC1 under visual context stimulation suggests that prediction-error detection engages a distinct dynamic, and the restriction of PC1 weights to anterior MOs raises the possibility of functionally relevant subdivisions within this area.

Reliable decoding required datasets free of pre-peak dilation and with a sufficiently large peak diameter; violating either criterion left performance indistinguishable from the null model. The first suggests that sustained and transient LC activities are mechanistically distinct modes of pupil modulation; the second reflects that our stimulation elicited dilation only probabilistically. Because cortical arousal state fluctuates on an infra-slow timescale below 0.2 Hz (Turner et al. 2023; Raut et al. 2025), evoked dilation may have occurred preferentially when stimulation coincided with a permissive phase of this oscillation, consistent with the frequency dependence of decoding.

### Model architecture and feature importance estimation

A GRU was selected because neural activity is time-series data and the feature window is short (1.5 s) (Ajioka et al. 2024). Multicollinearity among features, previously addressed by feature clustering (Jahedi et al. 2017), PCA-based transformation (Takagi et al. 2017) and Elastic Net regularization (Lee et al. 2025), was handled by ICA, which maximizes statistical independence with residual collinearity quantified by VIF, a general contribution to xAI in systems neuroscience.

## Limitations

The LC likely operates in heterogeneous states not captured by our models, and its tonic modulation of sensory cortex gain and prefrontal function (Arnsten 2009; Polack et al. 2013) may have influenced our results. Reflections from the gratings also required a low-pass filter on the pupil trace, reducing temporal precision. Critically, feature importance does not imply causality; optogenetic or chemogenetic manipulation is required to establish whether the identified signals modulate evoked pupil dilation (Sara and Bouret 2012; Yang et al. 2021). Finally, the spontaneous-dilation data came from a separate open-source dataset recorded in a different mouse line at a different sampling rate, so differences between evoked and spontaneous conditions may partly reflect dataset-related rather than biological factors.

## Conclusion

Using explainable machine learning, we found that evoked and spontaneous pupil dilation were preceded by distinct cortical dynamics, with evoked dilation involving a retrosplenial signal shared across sensory modalities and recruitment of the secondary motor cortex specific to visual stimulation. Reliable decoding requires the absence of pre-stimulus dilation, indicating that sustained and transient LC activity are distinct modes of pupil modulation. These findings provide a foundation for targeted investigation of cortical circuits linking sensory processing to arousal.

## Author contributions

T. K. and M. M. conceptualized the study. T. K. performed the experiments and preprocessed and analyzed the data. T. K., S. M., N. Y., O. M., N. N., T. T., S. T., and M. M. established the imaging setup and developed the analytical pipeline. T. K. and M. M. wrote the manuscript, and all the authors revised, read, and approved the final version of the manuscript.

## Data Availability

Data supporting the findings of this study are available from the corresponding author upon reasonable request.

## Code Availability

All custom codes used for cortical image analysis, pupil diameter quantification, and machine learning are publicly available at https://github.com/moritalabC201/WF_deep-learning.

## Supplementary materials

Supplementary Methods; Supplementary Fig. 1 to 12.

## Ethics statement

All animal procedures were approved by the Institutional Animal Care and Use Committee of Kobe University and conducted in accordance with institutional guidelines.

## Acknowledgements

The authors thank Ms. Yukiko Ohtaku for her assistance with mouse maintenance and technical support.

