## Supplementary materials for "Decoding Cortical Precursors of Evoked and Spontaneous Pupil Dilation"

### Supplementary Methods

#### Surgery and head-plate implantation.

Transgenic mice were housed with unrestricted access to food and water under controlled temperature (23 ± 2 °C), humidity (50 ± 10%), and a 12 h light/dark cycle. For transcranial optical clearing, mice were anesthetized by intramuscular injection of ketamine (100 mg/kg; Daiichi-Sankyo, Tokyo, Japan) and xylazine (10 mg/kg; Bayer Yakuhin, Osaka, Japan). The scalp was incised, and connective tissue on the skull surface was removed with a cotton swab. 4-META/MMA-TBB dental resin (Super-Bond C&B; Sun Medical, Moriyama, Japan) was applied to the skull surface, and a 3D-printed polylactic acid (PLA) frame (outer dimensions, 15 × 12 mm; frame width, 1 mm; thickness, 0.2 mm) with an 8 mm high light-shielding wall was secured to the edge of the skull. A 3D-printed plastic base (10 × 3 × 5 mm) and an aluminum headplate (10 × 40 × 1 mm) were then secured to the posterior skull with the same resin. Animals were monitored daily throughout the experimental period, and humane endpoints were defined as an abrupt loss of more than 20% of body weight within three days, or a loss exceeding 20–25% of the initial body weight. On completion of the experiments, mice were euthanized by cervical dislocation under deep pentobarbital anesthesia.

#### Optical configuration and image acquisition.

Mice were head-fixed under a custom-built fluorescence macroscope by securing the headplate with screws. TTL signals were generated by two pulse generators (SEN-7203, Nihon Kohden, Tokyo, Japan; Master-8, A.M.P.I., Jerusalem, Israel). GCaMP6f was excited using two LEDs combined through a dichroic mirror (DM435; 87-063, Edmund Optics, Barrington, NJ): for 480 nm excitation, a broadband white LED (SOLIS-1D; Thorlabs, Newton, NJ) with a bandpass filter (BP 480/40; D480/40m, Chroma Technology, Bellows Falls, VT); and for 405 nm excitation, a violet LED (SOLIS-405D; Thorlabs) with a bandpass filter (BP 405/10; Edmund Optics). Excitation and emission light were separated by a dichroic mirror (DM505; U-MNIBA, Evident, Tokyo, Japan), and fluorescence emission was further filtered (BP 530/40; U-MNIBA, Evident) before being detected by a scientific CMOS camera (ORCA-Flash 4.0 V3; Hamamatsu Photonics, Hamamatsu, Japan) controlled with HC Image software (Hamamatsu Photonics). Fluorescence was projected onto the camera sensor via a 2× objective lens (CFI Plan Apochromat Lambda D 2×; Nikon, Tokyo, Japan) and a 0.7× tube lens (focal length f = 70 mm, Thorlabs). Images were acquired as 16-bit TIFF files with a 15 ms exposure.

**Synchronization of pupil and cortical recordings.**

Video acquisition was initiated through a GPIO input on the Raspberry Pi. This approach synchronized the onset of recording rather than the acquisition timing of individual frames. Pupil images were acquired at 60 Hz (16.7-ms frame interval), but the accuracy of synchronization was limited by software-dependent latency and jitter in TTL detection and in the initiation of video recording.

**Registration and hemodynamic correction.**

Lambda and the medial and lateral edges of the olfactory bulbs were used as anatomical landmarks for affine registration (rotation and translation) to the cortical parcellation atlas. Pixel-wise fluorescence was normalized to the mean intensity measured during 5–15 s after recording onset ($F_0$) to yield $\Delta F/F$ images, where $\Delta F/F = (F - F_0)/F_0$. Singular value decomposition (SVD) was then applied to extract 34 spatial components and their corresponding temporal components. To separate the $Ca^{2+}$-dependent fluorescence signal from hemodynamic artifacts, the SVD temporal components were decomposed into $F_{405}$ and $F_{480}$ signals. Both signals were high-pass filtered at 0.1 Hz, a linear regression model was then fitted by ordinary least squares to predict $F_{480}$ from $F_{405}$, and the resulting estimate ($F'_{480}$) was subtracted from the measured $F_{480}$.

**Pupillometry hardware and pupil tracking.**

A near-infrared lamp (JC Infrared Illuminator; Xsdjasd, China) was used to illuminate the eye of the mouse, and an infrared camera (Raspberry Pi NoIR Camera V2; Raspberry Pi, Cambridge, UK), controlled by a Raspberry Pi 4 Model B (Raspberry Pi), was placed 5 cm in front of the right eye. The camera was controlled using custom Python code incorporating the Picamera library, and the acquired videos were saved in MP4 format. Pupil diameter was quantified using a custom Python 3.9 pipeline: the eye region was cropped from each frame, converted to grayscale, and downsampled to 20 Hz. To create the DeepLabCut training dataset, 90 images in total (30 images per mouse from three mice) were manually annotated. Labeled landmarks were the medial and lateral canthi (leye and reye) and nine points on the pupil: the center (center) and eight boundary points (top, bottom, left, right, tl, tr, bl, br). The network was trained for 5,000 iterations, and the resulting model was applied to all videos. Horizontal pupil diameter was converted to millimeters using a calibration image acquired under identical optical conditions. Frames in which any coordinate deviated more than 0.3 SD from the mean within a 0.5 s sliding window were identified as potential outliers, and the resulting traces were additionally inspected visually.

**Stimulation details.**

For somatosensory stimulation, two ECG electrodes (Red Dot Resting EKG Electrode 2360; 3M, St. Paul, MN), each cut to a width of 3 mm, were used, and stimulation was delivered through an isolator (ISO-Flex; A.M.P.I.). The threshold current was determined by increasing the current from 0.06 mA in 0.02 mA steps until pupil dilation was produced in at least 3 of 5 trials. Visual stimuli were generated using the PsychoPy library (https://www.psychopy.org/) and synchronized with data acquisition. In the visual context experiment, Redundant directions were chosen from 0°, 45°, 90°, and 135°, and the corresponding Deviant directions were 90°, 135°, 180°, and 225°, respectively. Each experiment consisted of eight sessions, with the Redundant direction randomly assigned as 0° or 45° in the first four sessions and as 90° or 135° in the last four.

**Resting-state preprocessing and labeling.**

Images of the open-source dataset were spatially downsampled to 288 × 288 pixels, and non-cortical pixels were masked to yield final images of 256 × 270 pixels. Resting-state pupil traces were low-pass filtered at 0.2 Hz and segmented into 40 s epochs from 50 to 570 s of the recording. The time window from −3 s to +0.5 s relative to the pupil diameter peak was Z-scored using the mean and standard deviation of the epoch, min−max normalized, and binarized at a threshold of 0.6.

**Cortical features based on Region of Interest (ROI).**

Hemodynamic-corrected GCaMP6f fluorescence was extracted from ROIs delineated based on 13 cortical regions in each hemisphere (26 regions in total) of the standard atlas (Supplementary Fig. 1), low-pass filtered at 2 Hz, Z-scored, and min-max normalized for use as input features. ROI-based features were used only for stimulation data.

**Independent component analysis.**

ICA was performed with the JADE implementation available at https://github.com/gbeckers/jadeR. Pixel-wise Z-scored $Ca^{2+}$ images were first decomposed by SVD, retaining the top 400 components. SVD temporal components for all sessions were obtained by projecting the data onto the SVD spatial basis and dividing by the corresponding singular values. ICA was applied to the top 200 SVD temporal components (cumulative explained variance > 99.5%) to yield whitened IC temporal components. ICA spatial components were reconstructed by right-multiplying the ICA mixing matrix by the SVD spatial basis.

**Dataset selection.**

Preliminary experiments with visual pattern stimulation indicated that the decoder failed to predict pupil dilation when the dataset included sessions in which the pupil reached the threshold before a specified pre-peak time window, or sessions in which the pupil never reached the threshold after stimulation (Fig. 2G; Supplementary Fig. 5B). Such sessions were therefore excluded from the dataset.

**Decoder implementation.**

All decoders were coded in-house (see Code Availability) and executed in Python 3.6. GRU models were implemented using the Keras API within TensorFlow and were trained using the Adam optimizer with binary cross-entropy loss and a mini-batch size of 256. LSTM models replaced the GRU layer with a single LSTM layer (32 units), retaining the same fully connected output structure, sigmoid activation, optimizer, loss and mini-batch size. For Linear SVM models, IC temporal feature matrices were flattened into a single vector per sample, and models were trained by minimizing the squared hinge loss for 15,000 iterations. Classification accuracy was obtained by binarizing model predictions at a fixed threshold of 0.5 and computing the proportion of correctly classified samples; the accuracy at the epoch with the lowest validation loss was used as the summary metric for each model.

**Multicollinearity and feature importance.**

Validation data were used to estimate feature importance. Prior to the estimation, multicollinearity among features within each time frame was quantified as the variance inflation factor ($VIF = 1/(1 - R^2)$, where $R^2$ is the coefficient of determination obtained by regressing each feature on all others), and models with maximum $VIF < 10$ were subjected to the estimation. The importance of each feature was quantified as the decrease in ROC-AUC following random permutation of that feature's values across samples. Random permutation was repeated three times, and the mean was taken as the final estimate.

**Deep SHAP implementation.**

Deep SHAP was applied using the SHAP Python package (https://github.com/slundberg/shap). Mapped SHAP value arrays had dimensions $n \times T \times H \times W$ (n, number of models; T, number of time frames; $H \times W$, image resolution) for each stimulation condition. PCA temporal components for null models and resting-state data were obtained by projecting the corresponding mapped SHAP values onto the PCA spatial components and dividing by the respective singular values.

**Choice of statistical tests.**

Paired t-tests were used to assess differences between matched observations, namely between trained decoders and their corresponding null decoders, between SHAP-based measures, and between measurements obtained under different stimulation conditions. Wilcoxon signed-rank tests were used for nonparametric comparisons between paired observations, whereas Mann–Whitney U tests were used for comparisons between independent groups. The assumption of normally distributed paired differences underlying the paired t-tests was not verified.

**Supplementary Figures**

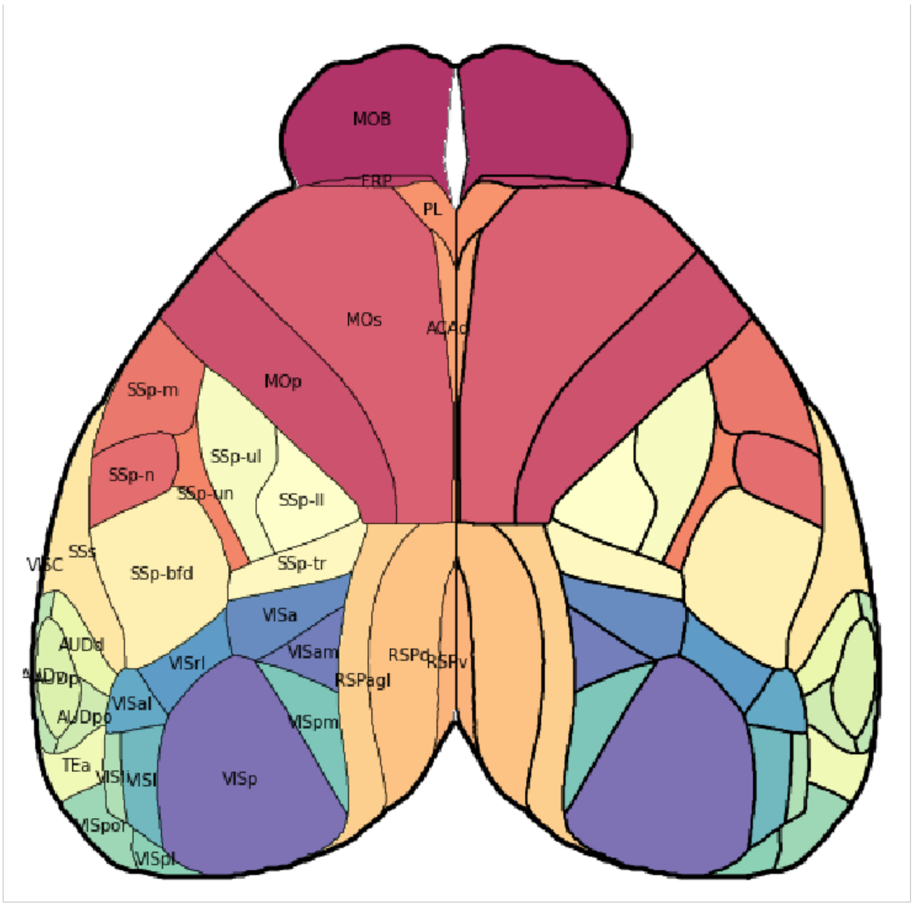


VISp, primary visual area,
VISam, anteromedial visual area,
VISa, anterior area,
VISpm, posteromedial visual area,
SSp-ul, primary somatosensory area-upper limb,
SSp-ll, primary somatosensory area-lower limb,
SSp-tr, primary somatosensory area-trunk,
RSPagl, retrosplenial area-lateral agranular part,
RSPd, retrosplenial area-dorsal part,
RSPv, retrosplenial area-ventral part,
ACAd, anterior cingulate area-dorsal part,
MOs, secondary motor area,
MOp, primary motor area

**Supplementary Figure 1. Brain region nomenclature in the standard brain atlas.**

Brain region names follow the nomenclature of the Allen Mouse Brain Atlas (Allen Institute for Brain Science (2011). Allen Mouse Brain Atlas. Available from: https://atlas.brain-map.org). VISp, Primary visual area; VISam, Anteromedial visual area; VISa, Anterior area; VISpm, Posteromedial visual area; SSp-ul, Primary somatosensory area–upper limb; SSp-ll, Primary somatosensory area–lower limb; SSp-tr, Primary somatosensory area–trunk; RSPagl, Retrosplenial area–lateral agranular part; RSPd, Retrosplenial area–dorsal part; RSPv, Retrosplenial area–ventral part; ACAd, Anterior cingulate area–dorsal part; MOs, Secondary motor area; MOp, Primary motor area.

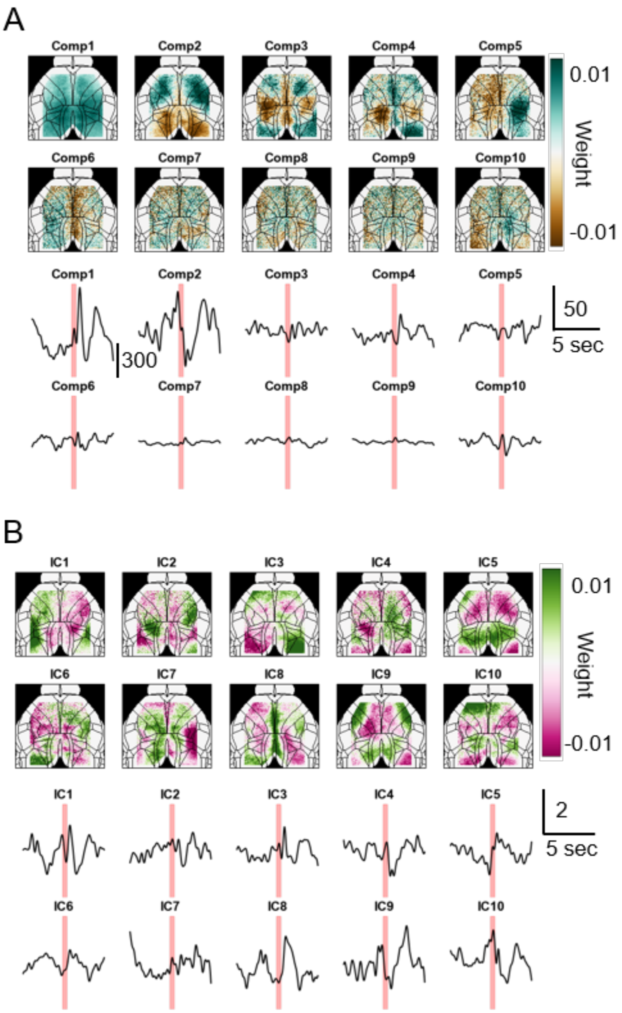

**Supplementary Figure 2. Feature extraction of somatosensory-evoked cortical activity.**

***A,*** Representative (Top 10) spatial (upper) and temporal (lower) components (Comp1–Comp10) extracted from cortical activity images of the somatosensory stimulation dataset (N = 3 mice, n = 23 sessions; every fifth session) by SVD. ***B,*** All the spatial (upper) and temporal (lower) independent components (IC1–IC10) obtained by applying ICA to 200 SVD-extracted components.

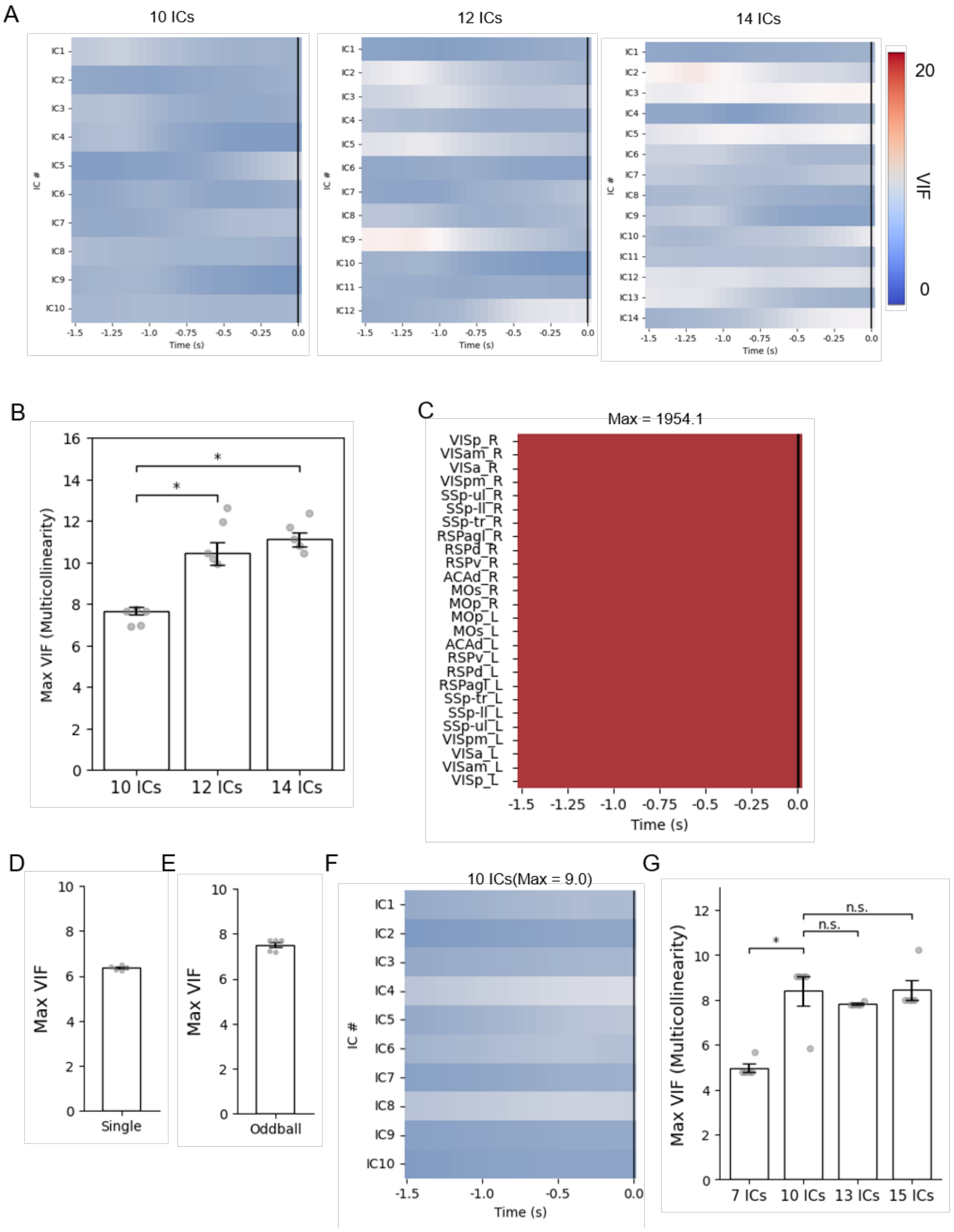


**Supplementary Figure 3. Assessment of the effect of feature selection on multicollinearity by the variance inflation factor (VIF).**

***A*****,** Model-averaged heatmaps showing the time-dependent change of VIF before somatosensory stimulation onset (0.0 s), when 10, 12, and 14 ICs were used as features. ***B,*** Maximum VIF for each number of input ICs. Median ± SE (n = 5 models). Mann–Whitney U test with Holm correction; *P < 0.05. ***C,*** Model-averaged heatmap showing the time-dependent change of VIF when ROIs were used as features. ***D,*** Maximum VIF for visual pattern stimulation. ***E,*** Maximum VIF for visual context stimulation. ***F,*** Representative heatmap showing the time course of VIF before spontaneous pupil dilation onset, when 10 ICs were used as features. ***G,*** Maximum VIF when 7, 10, 13, or 15 ICs were used as features. Median ± SE. Mann–Whitney U test with Holm correction; n.s. P > 0.05.

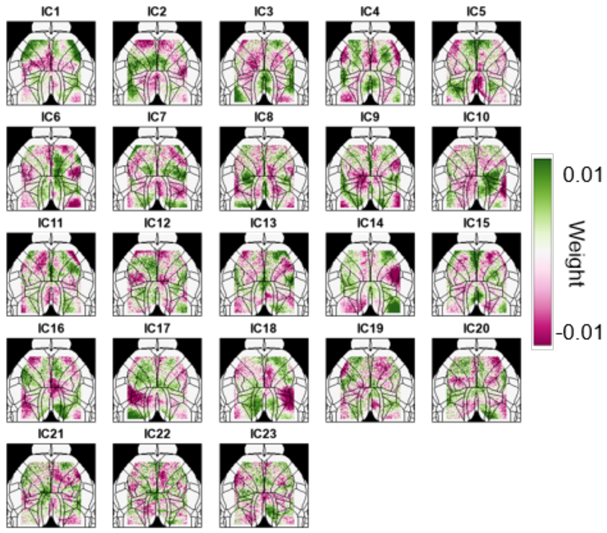


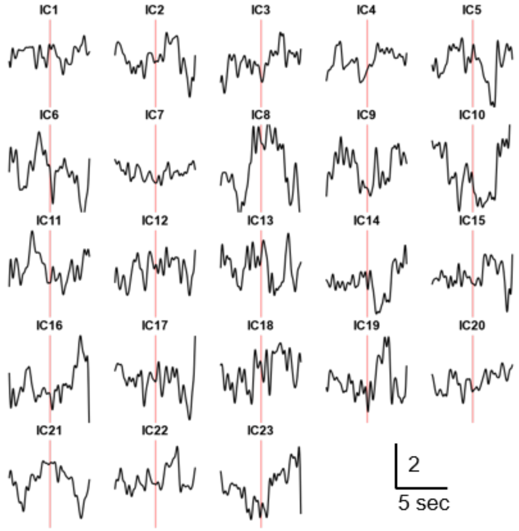

**Supplementary Figure 4. Feature extraction of cortical activity evoked by visual pattern stimulation and subsequent preprocessing for decoder training.**

Spatial (upper) and temporal (lower) independent components (IC1–IC23) obtained by applying ICA to 200 SVD-extracted components from all datasets with visual pattern stimulation.

**Supplementary Figure 5. Optimization of pupil label threshold.**

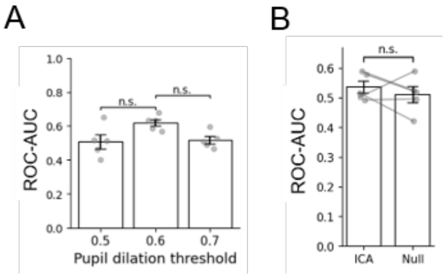


***A,*** Effect of pupil label threshold on model performance assessed by ROC-AUC. Mann–Whitney U test with Holm correction; n.s. P > 0.05. ***B,*** Comparison of ROC-AUC between RNN decoders trained with datasets below the post-stimulus threshold and those trained with null datasets. Paired t-test. n.s. P > 0.05.

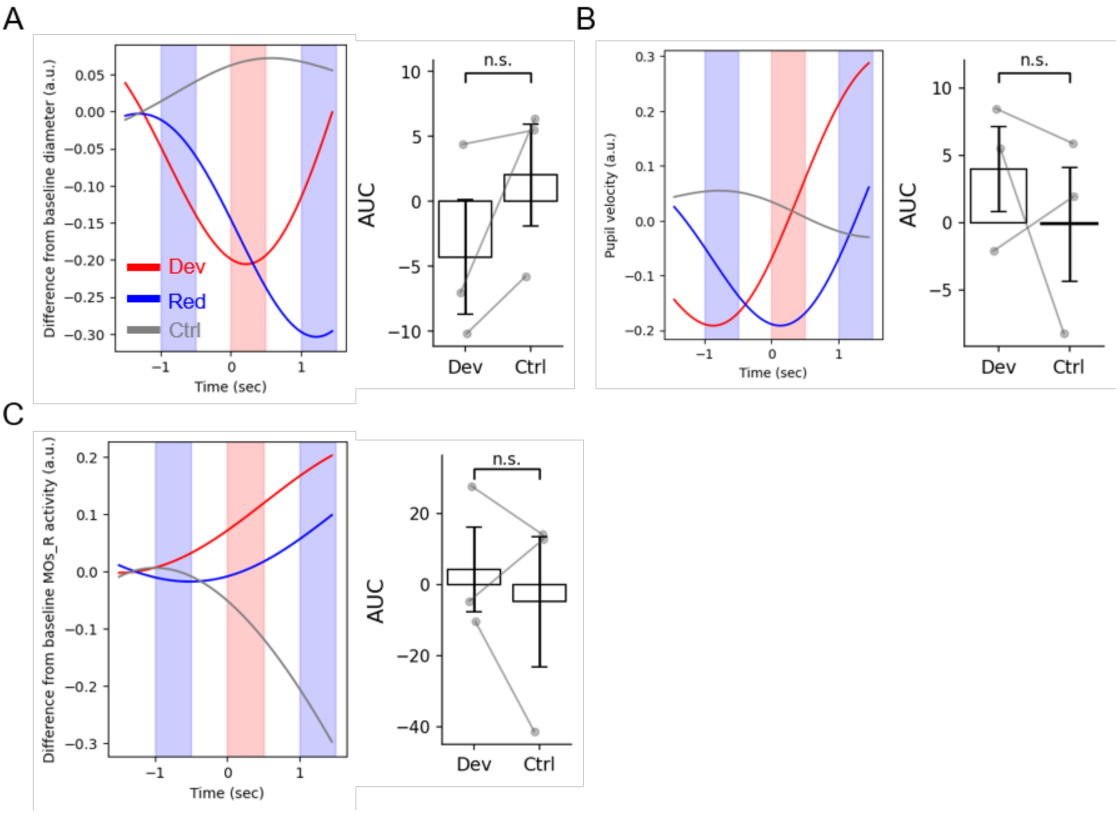

**Supplementary Figure 6. Pupil dilation and cortical activity in response to visual oddball paradigm.**

***A,*** *Left,* Mean traces of pupil diameter deviation from baseline. Deviant stimulation (Dev); Redundant stimulation (Red); Random control stimulation (Ctrl). *Right,* Area under the curve of pupil diameter traces, AUC (pupil). ***B,*** *Left,* Mean traces of pupil dilation velocity. *Right,* Area under the curve of pupil dilation velocity traces, AUC (pupil velocity). ***C,*** Mean traces of cortical activity in right secondary motor area (MOs_R) 1.5 s before and after stimulus onset (left). Area under the curve of cortical activity traces, AUC (MOs_R activity) (right). Mean ± SE. Paired t-test. N = 3. n.s. $P > 0.05$ for A, B and C.

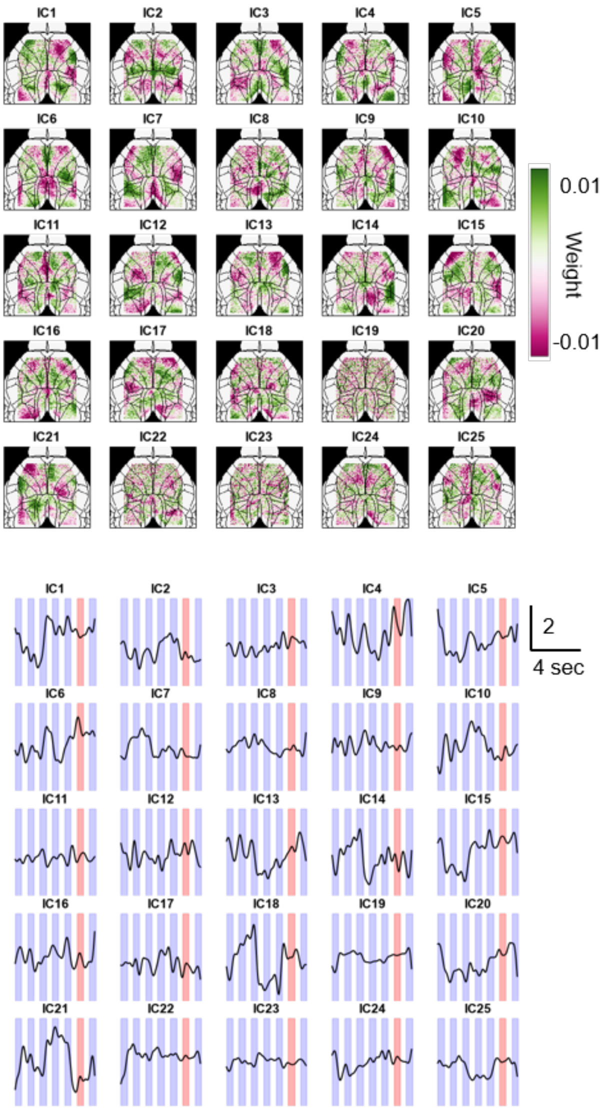


**Supplementary Figure 7. Feature extraction of cortical activity evoked by visual context stimulation.**

Spatial (upper) and temporal (lower) independent components (IC1–IC25) obtained by applying ICA to 200 SVD-extracted components from all datasets with visual context stimulation. Blue and red shading indicate redundant and deviant stimulation periods, respectively.

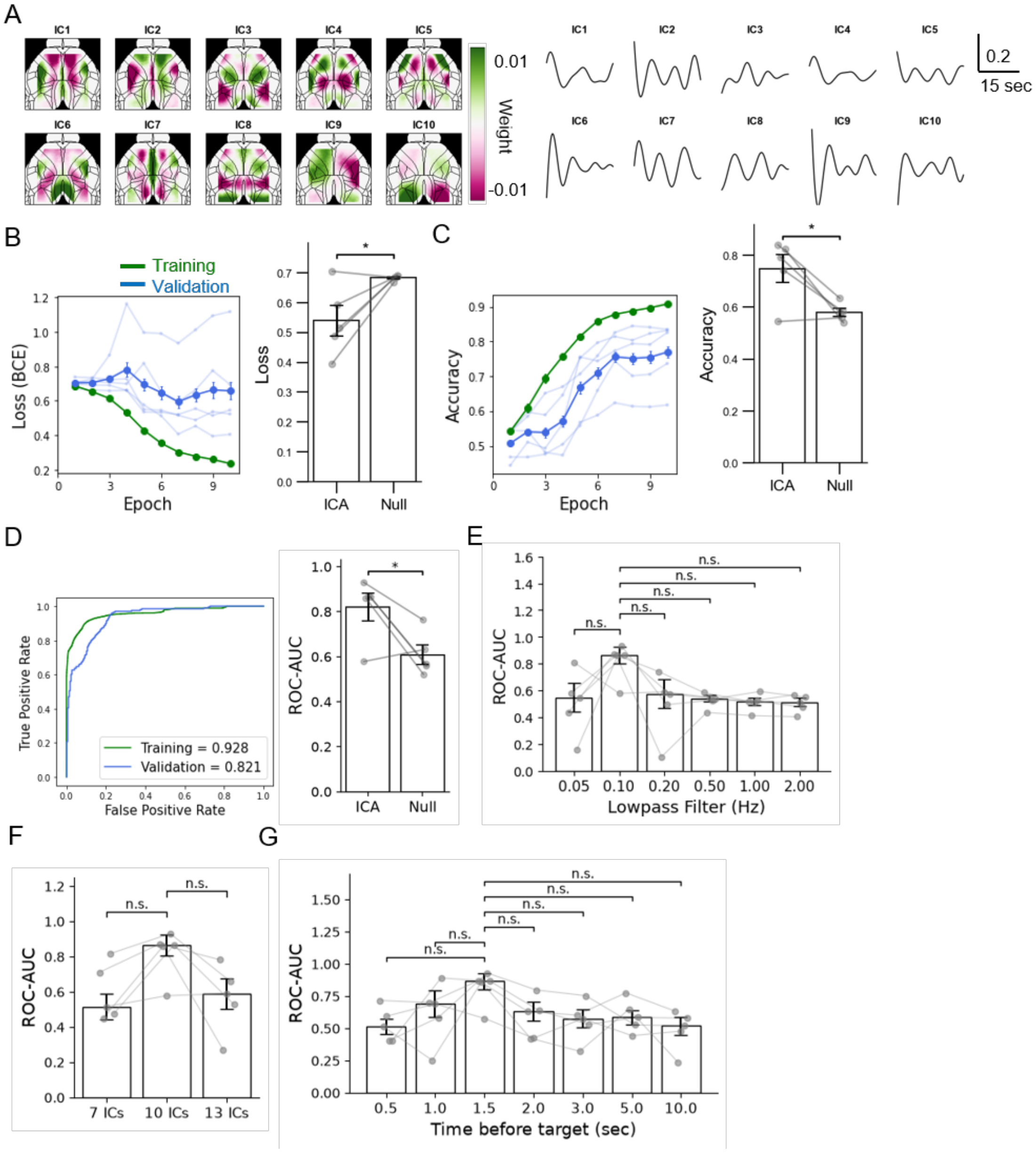


**Supplementary Figure 8. RNN decoder for spontaneous pupil dilation.**

***A,*** Spatial (left) and temporal (right) independent components (IC1–IC10) obtained by applying ICA to 200 SVD temporal components extracted from resting state data. ***B,*** Prediction by RNN-based decoder assessed by loss (BCE). Loss changes of training (Training) and validation (Validation) datasets over 10 epochs (Epoch) (*left*) and comparison with a null model trained on within-session target-shuffled datasets (*right*). The dataset was randomly divided into five equal folds; four were used for training and one for validation using a leave-one-out approach, generating models (n = 5). ***C,*** Prediction assessed by accuracy. ***D,*** Prediction assessed by ROC curves. ROC curves from a representative decoder, and the mean ROC-AUC of training and validation datasets across five models (*left*). Comparison of ROC-AUC against the null model (*right*). Mean ± SE (n = 5 models). Paired t-test; *P < 0.05 for B, C, and D. ***E,*** Effect of low-pass filter applied to ICs on model performance. ***F,*** Effect of the number of input ICs on model performance. ***G,*** Effect of start time of the ICA temporal component input window before target (Time before target) on model performance. Model performance was assessed by ROC-AUC for E, F, and G. Median ± SE (n = 5 models). Wilcoxon signed-rank with Holm correction; n.s. P > 0.05.

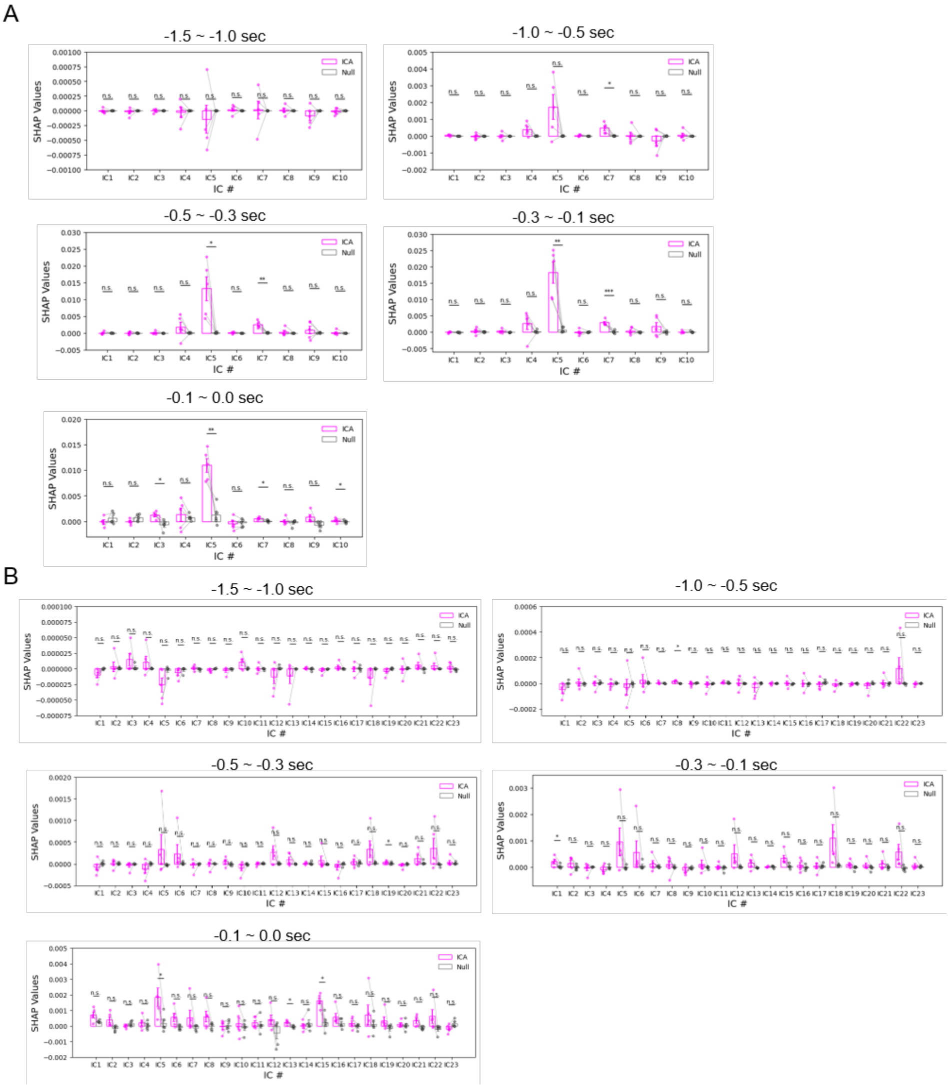

C

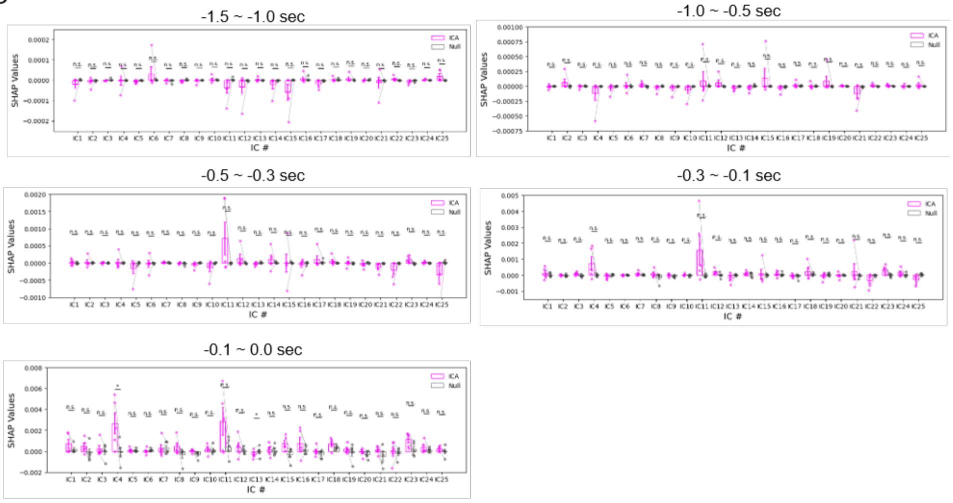


D

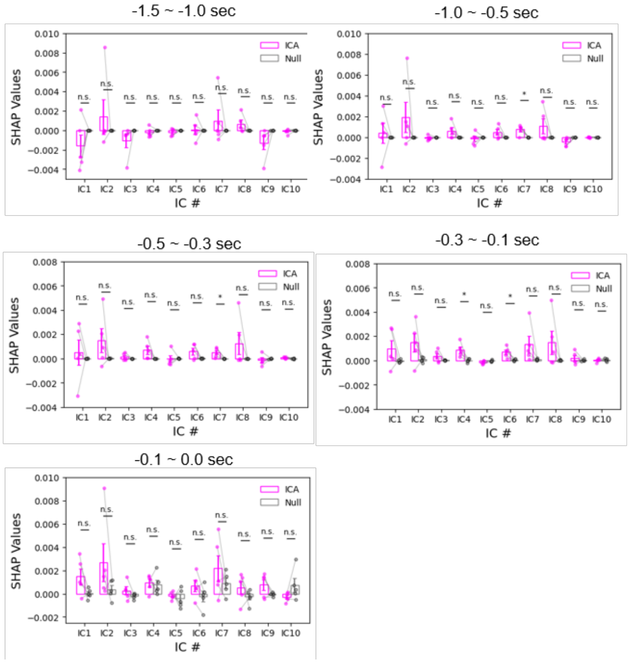


**Supplementary Figure 9. Statistical analysis of SHAP values.**

SHAP values of ***A*,** somatosensory, ***B***, visual pattern, ***C*,** visual context stimulation, and ***D*,** spontaneous pupil dilation decoders compared against null decoders during the indicated time windows. Mean ± SE (n = 5). Paired t-test (uncorrected); n.s. P > 0.05, *P < 0.05, **P < 0.01, ***P < 0.005.

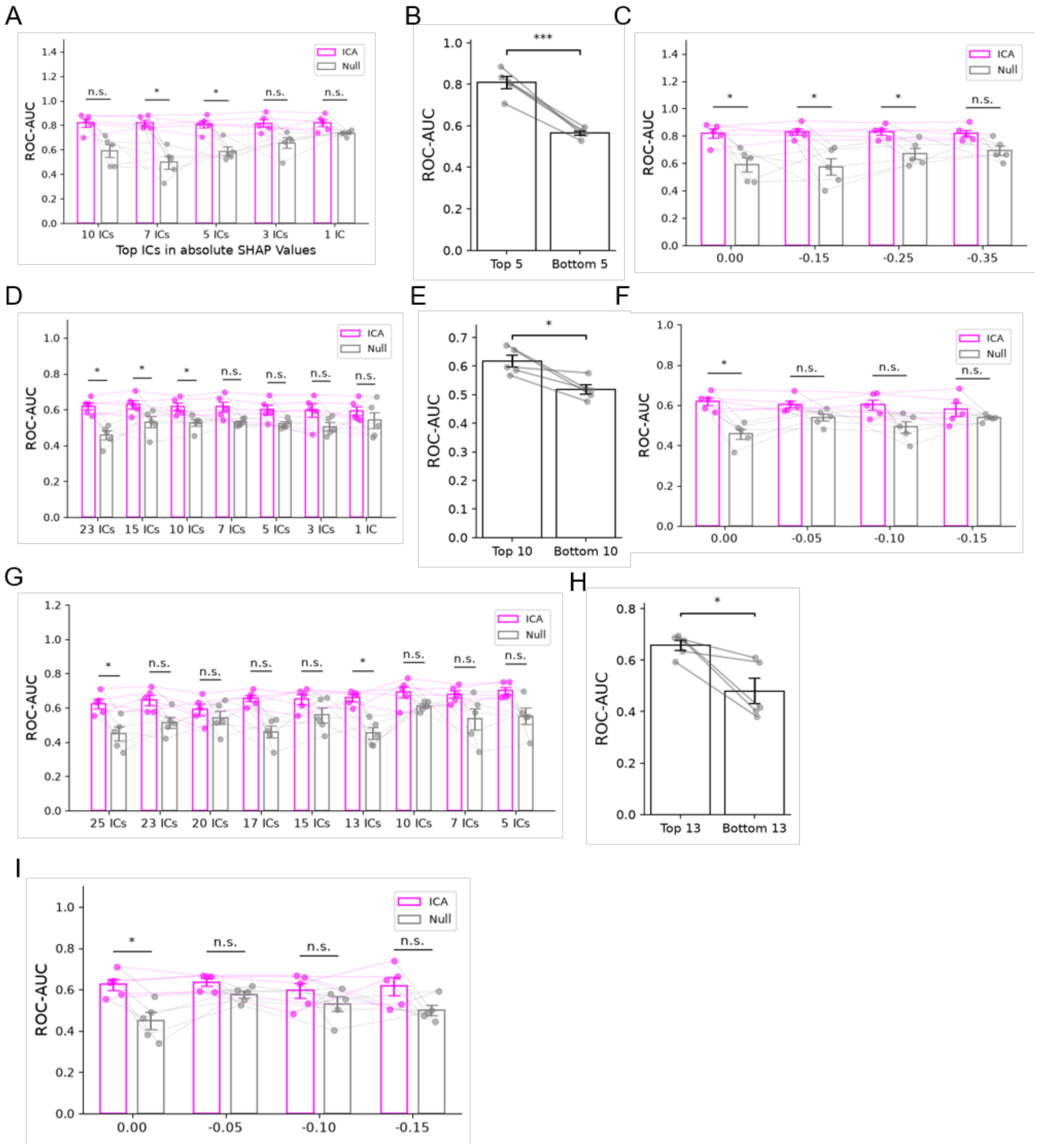


**Supplementary Figure 10. Validation of SHAP values by RNN-based decoder using absolute SHAP values.**

***A–C***, somatosensory stimulation. ***D–F,*** visual pattern stimulation. ***G–I,*** visual context stimulation***. A, D, G,*** AUC of decoders using ICs with high absolute SHAP values. Paired t-test with Holm correction. Mean ± SE (n = 5). n.s. P > 0.05, *P < 0.05. ***B, E, H,*** AUC for decoders using the top-ranked ICs (top 5, 10, or 13) as indicated by SHAP value, and the same number of bottom-ranked ICs. Mean ± SE (n

= 5). Paired t-test. *P < 0.05, ***P < 0.005. ***C, F, I,*** Comparison of AUC as a function of the end time of the input time window relative to the pupil-labeled frame. Paired t-test with Holm correction. Mean ± SE (n = 5). n.s. P > 0.05, *P < 0.05.

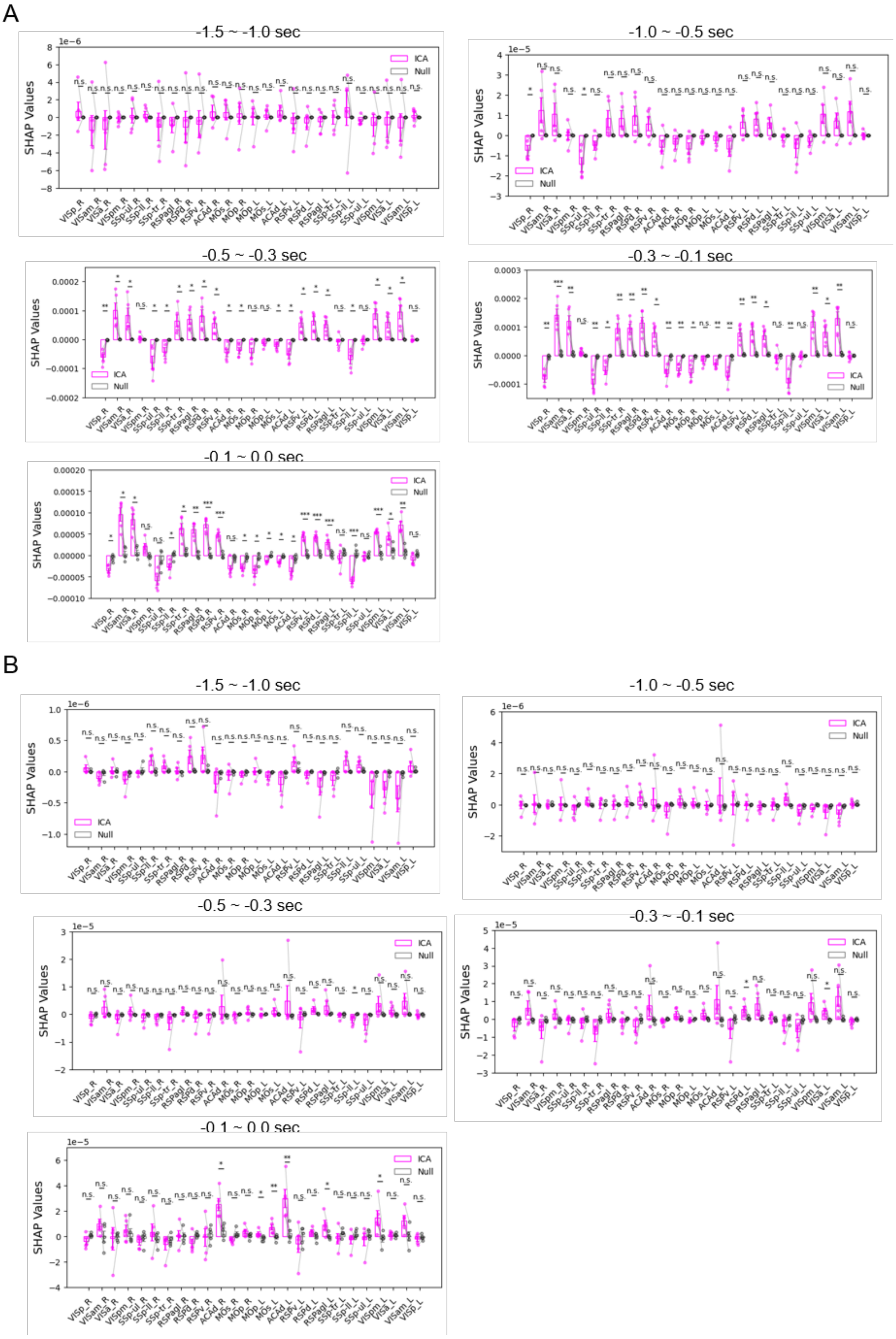

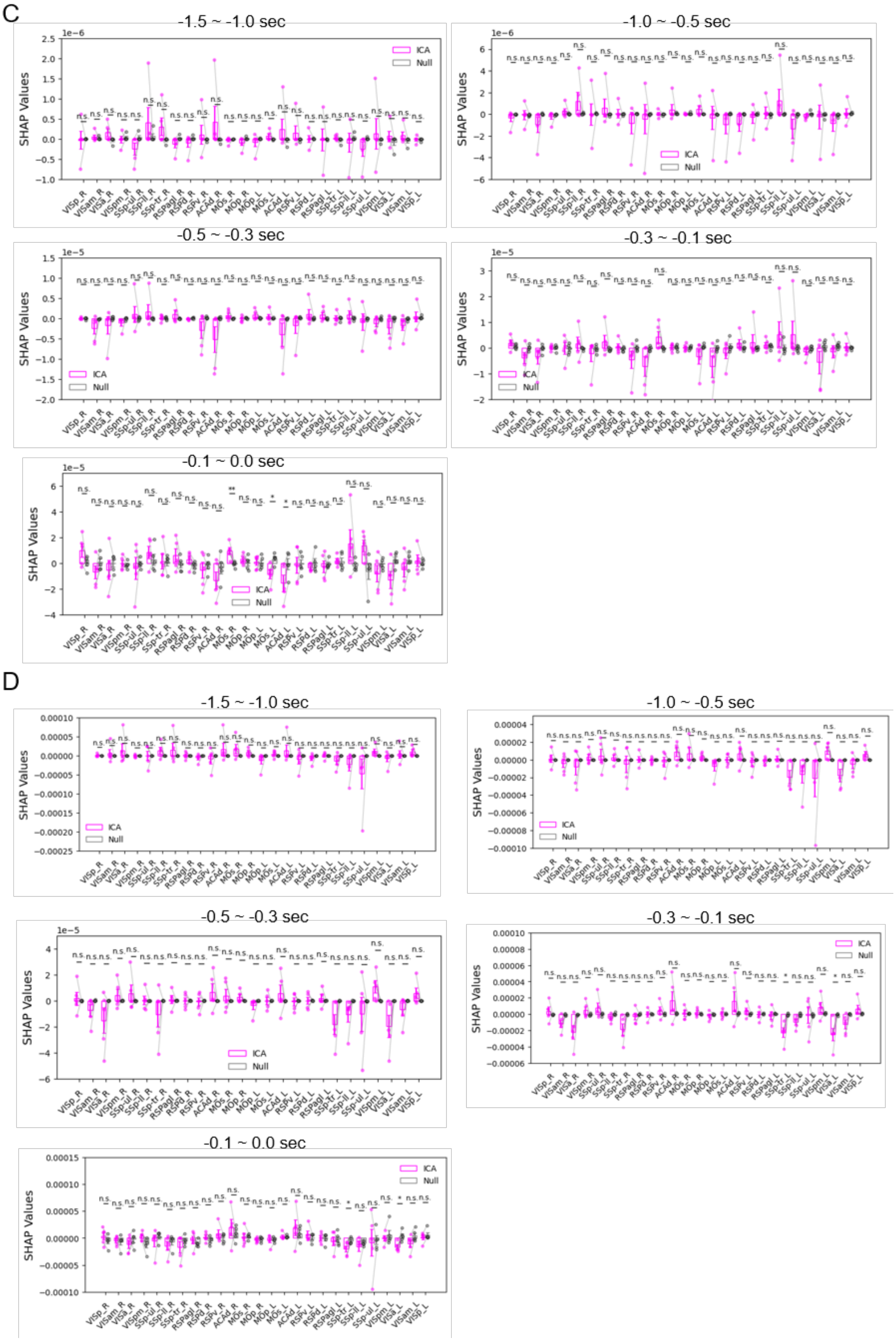


**Supplementary Figure 11. Mapping SHAP values based on brain regions.**

Mapped SHAP values during the indicated time window were compared with null decoders in ***A***, somatosensory, ***B,*** visual pattern, ***C,*** visual context stimulation, and ***D,*** spontaneous pupil dilation. Mean ± SE (n = 5). Paired t-test (uncorrected); n.s. P > 0.05, *P < 0.05, **P < 0.01, ***P < 0.005.

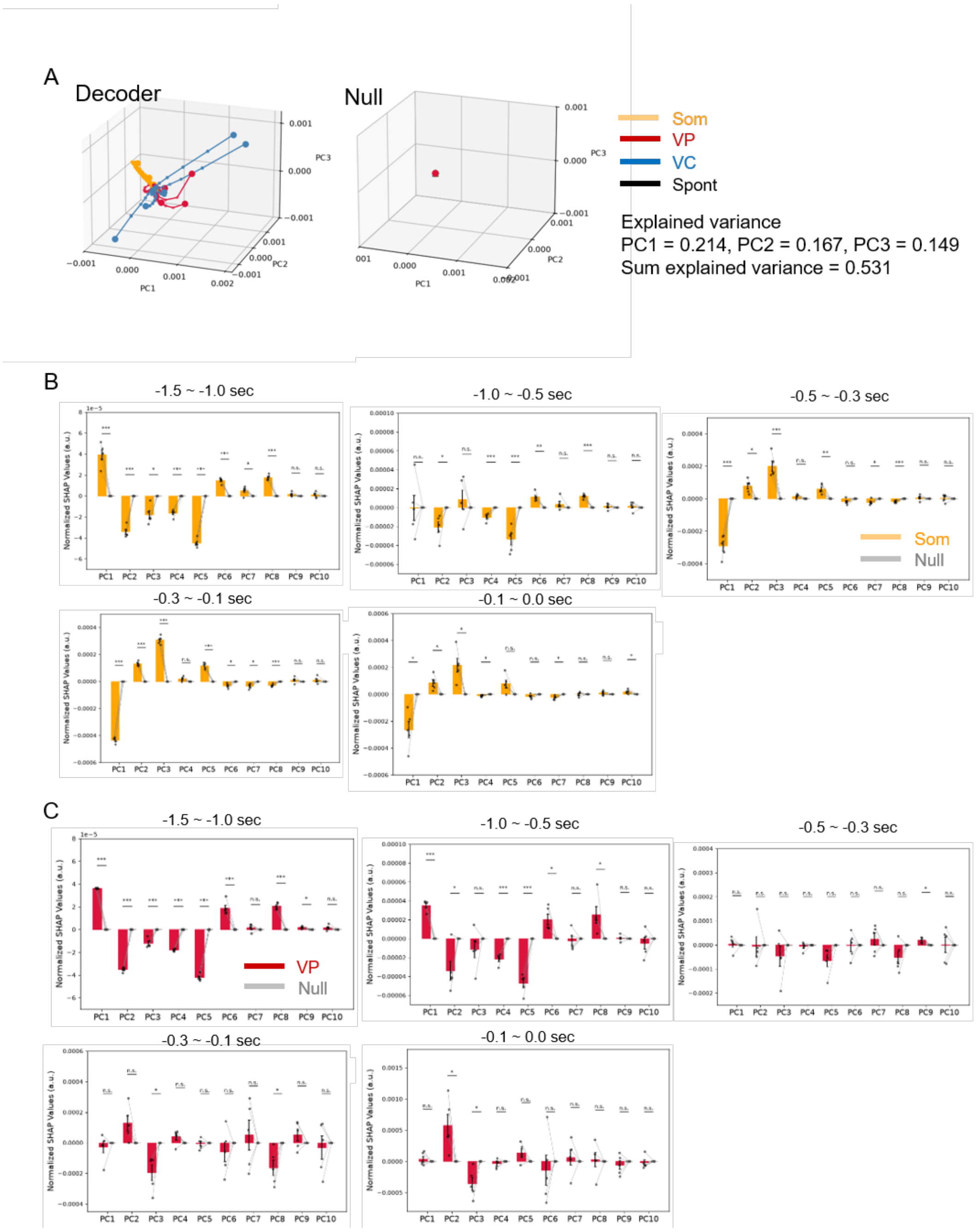

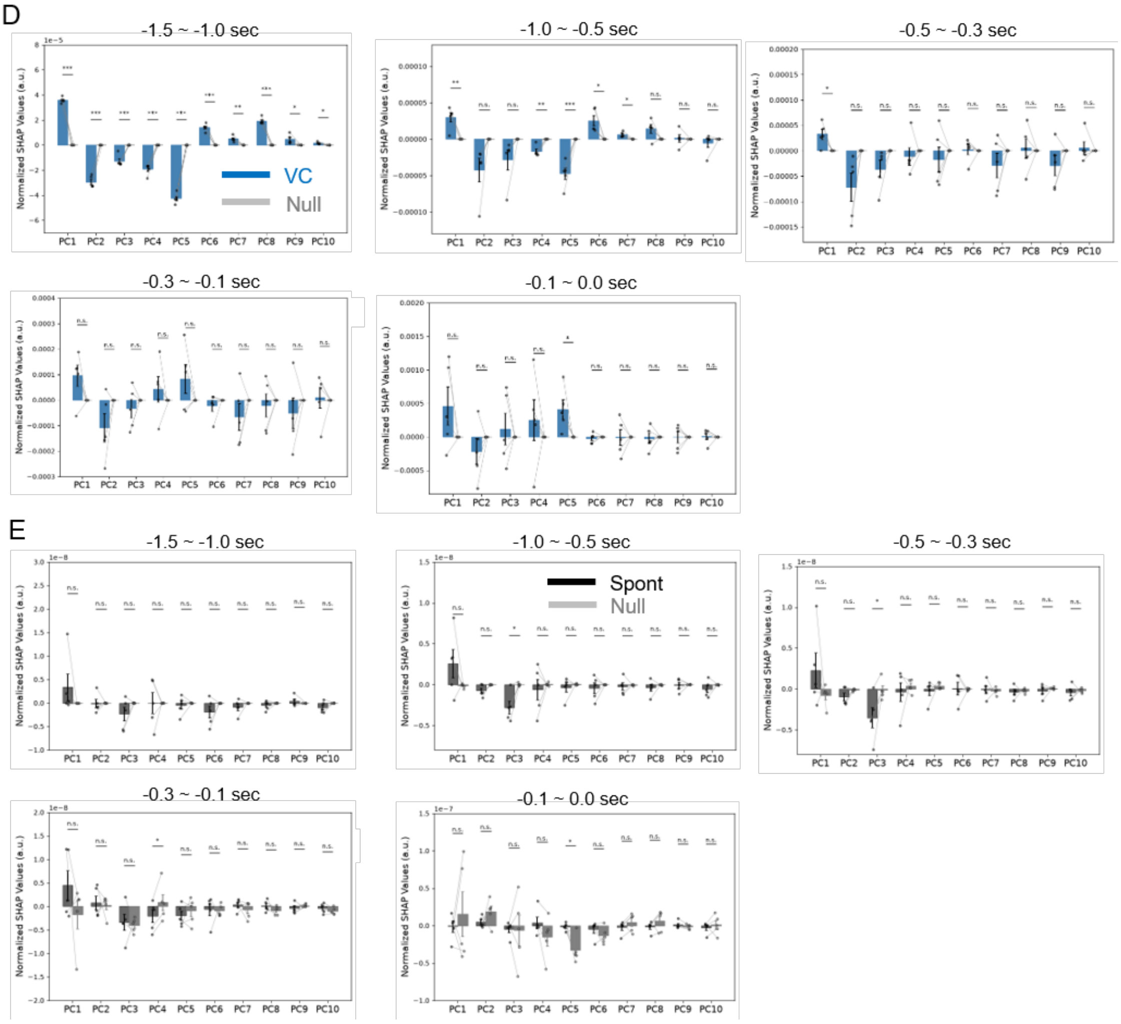


**Supplementary Figure 12. Presentation of feature contribution by principal components.**

***A,*** Trajectories of SHAP values were plotted in PC1–PC3 space. Decoders (*left*) and null decoders (*right*) for somatosensory stimulation (Som), visual pattern stimulation (VP), visual context stimulation (VC), and spontaneous dilation (Spont). Start and end points are indicated by triangles and circles, respectively. Temporal resolution for Som, VP, and VC is 0.05 s, while that for Spont is 0.03 s. The explained variance of each component and their cumulative value are indicated above the plots. Normalized SHAP values of PCA temporal components compared against null decoders during the indicated time windows for ***B***, Som, ***C,*** VP, ***D,*** VC, ***E,*** Spont. Mean ± SE (n = 5). Paired t-test; n.s. P > 0.05, *P < 0.05, **P < 0.01, ***P < 0.005.
